# A neural model of working memory

**DOI:** 10.1101/233007

**Authors:** Sanjay G Manohar, Nahid Zokaei, Sean J Fallon, Tim Vogels, Masud Husain

## Abstract

Working memory, the ability to keep recently encountered information available for immediate processing, has been proposed to rely on two mechanisms that appear difficult to reconcile: self-sustained neural firing, or the opposite—activity-silent synaptic traces. Here we show that both phenomena can co-exist within a unified system in which neurons hold information in both activity and synapses. Rapid plasticity in flexibly-coding neurons allows features to be bound together into objects, with an important emergent property being the focus of attention. One memory item is held by persistent activity in an attended or “focused” state, and is thus remembered better than other items. Other, previously attended items can remain in memory but in the background, encoded in activity-silent synaptic traces. This dual functional architecture provides a unified common mechanism accounting for a diverse range of perplexing attention and memory effects that have been hitherto difficult to explain in a single theoretical framework.

## Introduction

Our capacity to hold and manipulate information over delays of a few seconds has long been thought to be subserved by the persistent firing of neurons during the delay (Funahashi, 2017; Fuster and Alexander, 1971). However, a number of recent studies have instead proposed “activity-silent” working memory, in which synaptic weights hold information during the delay, even in the absence of neuronal firing (Silvanto, 2017; Sreenivasan et al., 2014; Mongillo et al., 2008; Stokes, 2015). This dispute comes at a time when it is also becoming clear that working memory (WM) is not a homogeneous store. When we hold multiple items in WM, strong attentional effects are apparent. For example, people are faster and more accurate to recall the last item encoded, or the last item that was brought to mind (Chun et al., 2011; Oberauer, 2002; Souza and Oberauer, 2016; Zokaei et al., 2014a).

One item in memory, termed the ‘focus of attention’, appears to be in a privileged state. It is decodable using functional MRI and is susceptible to TMS, unlike the unfocused items which considered to be stored but in a non-privileged state (Lewis-Peacock et al., 2012; Sprague et al., 2016). In contrast, unfocused items can only be decoded after their latent representation is reactivated (Rose et al., 2016; Wolff et al., 2017). Thus both active and inactive representations may coexist in WM, and items can move between these two states (LaRocque et al., 2014; Postle, 2016; Zokaei et al., 2014b). Computational neural models of both active (Compte et al., 2000; Zenke et al., 2015) and silent (Mi et al., 2017; Mongillo et al., 2008) WM have been separately postulated, but neither type of model on their own accounted for shifts of attention within WM. Here we unify these divergent approaches in a new class of memory model.

Rapid synaptic plasticity at the millisecond scale has been used to explain how a pattern of inputs can be remembered (Fiebig and Lansner, 2017; Sandberg et al., 2003). In these synaptic models, simultaneously-activated neurons become more strongly connected, so that when a partial pattern is later presented, the original combination of active neurons can be re-activated, by associative recall. At longer time scales, synaptic plasticity may underlie associative episodic memory (Burgess and Hitch, 2005; Rizzuto and Kahana, 2001), or long term memory, which can provide the synaptic backdrop to support an active WM (Litwin-Kumar and Doiron, 2014, 2014; Zenke et al., 2015). Rapid plasticity in auto-associative networks can also account for serial recall of sequences of items (Fiebig and Lansner, 2017; Howard and Kahana, 2002) – including serial order effects such as primacy and recency (Farrell and Lewandowsky, 2002) – because new information may use up free space, or overwrite old information (Matthey et al., 2015; Sandberg et al., 2003). Because the physiological meaning of a neuron’s firing depends upon its input and output connections, plasticity in these models could lead to neurons whose activity represents different things on different trials – a property that we characterize here as *flexible coding.* However these models do not produce stable persistent-activity states in feature-selective neurons, which has long been considered a hallmark of WM (Funahashi, 2017).

In contrast, in sustained activity models, items are held WM by virtue of delay-period activity (Compte et al., 2000; Funahashi, 2015; Funahashi et al., 1989), which relies on positive feedback to allow stimulus-induced activity to persist or resonate, leading to an “attractor” state. (Chumbley et al., 2008; Wimmer et al., 2014; Zipser et al., 1993). Although such active maintenance may also depend upon rapid changes in synaptic weights (Hansel and Mato, 2013; Pereira and Wang, 2015), these models do not generally allow memory recall from a silent inactive state.

The present work unites persistent activity attractors with silent synaptic storage. In our new class of memory model, both active and silent representations are essential to WM. We propose that persistent activation serves as the *focus of attention* that encodes recent activity patterns into synapses. Rapid plasticity in flexibly-coding neurons allows features to be bound together into objects, with an emergent property being that the last item is maintained actively. Recent, previously-attended items are preserved instead in synaptic traces. They are in a non-privileged state but, importantly, can be re-activated by partial information.

We propose that attention arises from the interaction between two distinct types of neural representation: fixed *feature* neurons, and *freely-conjunctive* neurons (**Fig.1A**). Feature neurons may be sensory, motor or conceptual. They have fixed receptive fields or tuning curves – as observed in posterior cortical areas. In contrast, the freely-conjunctive neurons can rapidly change their connection weights with the feature cells, and therefore their activity does *not* represent a fixed feature or item in memory. Instead, through rapid plasticity on each trial, a conjunctive cell will come to encode a conjunction of simultaneously active features, by forming a reciprocal associative mapping to feature-selective neurons.

**Fig.1:**
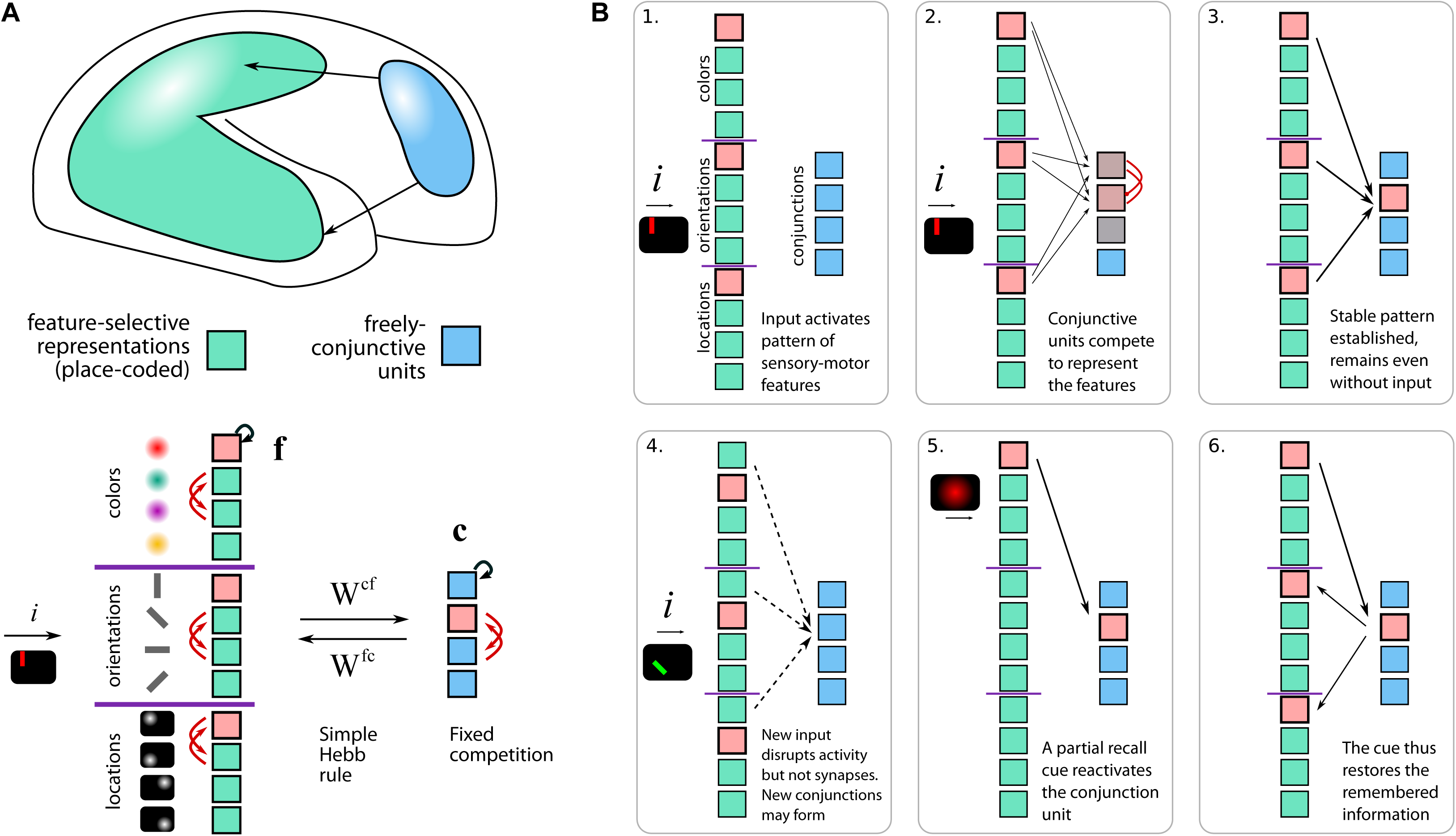
Conjunctive neurons to support attention and working memory. **A** Two populations of neurons are distinguished based on their inputs. Posterior neurons (green) encode sensory-motor features, whereas prefrontal neurons (blue) are “conjunctive”: i.e. they are able to rapidly increase or decrease their synaptic connectivity with patterns of feature neurons, using a Hebbian associative rule. We simulated 12 feature-selective neurons (**f**) and 4 freely-conjunctive neurons (**c**). An active combination of neurons (pink) causes strengthening of synapses in both directions, producing a stable attractor across brain areas. **c**=conjunctive cells, **f**=feature cells. W=synaptic weights, i=sensory input. **B** Sequence of proposed neuronal events during attention, encoding and retrieval in working memory. **1.** Sensory input activates features. In this case a vertical red bar located at the top left of the display activates separate feature neurons tuned to orientation, color and location. **2.** Features excite conjunctive neurons, which compete. **3.** The winning conjunction drives sustained activity. **4.** New input to the system (in this case an oblique purple bar at bottom left) disrupts current firing activity, but synaptic weightings remain. **5.** Probe feature (in this case red colour) re-activates the original conjunctive unit that encoded the red vertical bar. **6.** Conjunctive unit re-activates original features, completing recall.

Persistent activity arises by mutual excitation between feature and conjunction neurons. The conjunction neurons form a limited-capacity store that can hold many kinds of information in one place. Thus, our model bridges the gap between neuron-level descriptions and the psychological notion of a *general-purpose register,* sometimes termed a “memory slot” (Cowan, 2010; Luck and Vogel, 1997), a concept which has not as yet been characterized at the level of single prefrontal neurons. Such registers are difficult to explain unless individual neurons can encode different types of WM content at different times. Our model permits this by allowing rapid synaptic changes so conjunctive neurons can represent many kinds of information, depending on the recent context.

We suggest that two lines of evidence point to such conjunction neurons being located in prefrontal cortex (PFC): firstly, PFC is highly active in memory and manipulation (Eriksson et al., 2015; Postle et al., 2006), yet secondly, information is not always easy to decode (Christophel et al., 2012; Cogan et al., 2017; Kammski et al., 2017). Although WM contents can undoubtedly be decoded from many PFC neurons, about 60% of prefrontal neurons appear to be nonselective, and even for those that are selective, they often show less than a 50% modulation of their firing rate by information in WM (Miller et al., 1996; Parthasarathy et al., 2017). This apparently-nonselective component of prefrontal activity could reflect transient and flexible coding by conjunctive units.

In this study, we first aim to provide a single common mechanism accounting for a diverse range of perplexing attention and memory effects. Second, we attempt to explain neurophysiological data where items in memory initially produce persistent activity, which then falls “silent” when attention shifts to new information (Konecky et al., 2017). Third, we aim to explain why many imaging studies conclude that attention and working memory are “distributed” processes involving both prefrontal and sensory brain areas (Christophel et al., 2017; Gayet et al., 2017, 2017; Xu, 2017). In our simulations, we chose to examine the extreme situation where conjunctive neurons are fully nonselective for features. This limiting scenario is of course implausible, since no single prefrontal neuron could receive input from every feature neuron. However we argue that it is a highly illustrative paradigmatic case. In reality prefrontal neurons will necessarily have some degree of selectivity, but here we focus only on characterizing the novel concept of how rapid plasticity can give rise to flexible coding, and therefore we model *purely* conjunctive neurons as distinct from feature-selective neurons.

## Results

### 1. Operation of the Model

When a stimulus is perceived (**Fig.1B; Movie S1**), conjunctive neurons compete through lateral inhibition to become active in response to the combination of active features. In the example shown in **Fig.1** the conjuncton units learn rapidly to encode combinations of color, orientation and location (**Fig.1B.2**). During encoding into WM, the winning conjunctive unit sustains the activity of all co-active feature neurons through mutual excitation. This strengthens synapses in both directions through rapid Hebbian plasticity, further stabilizing the active pattern. Once a conjunctive unit succeeds in reciprocally activating a set of feature units, *attention is focused* on the activated features, binding the features of a compound stimulus into a perceptual object.

The reciprocal feature-to-conjunctive synapses keep the novel combination of features persistently active, even when the stimulus is no longer present (**Fig.1B.3**).

When a new stimulus arrives, a new pattern of sensory input destabilizes internal activity, thus triggering a shift of attention towards the newly activated features. A new conjunction may win, to become the new focus of attention. Crucially, however, synapses between the previous object’s constituent features and one particular conjunctive unit remain strengthened even after those neurons fall silent (**Fig.1B.4**). Thus, presenting any one feature of a previously attended object (e.g. color, as shown in **Fig.1**) will act as a memory probe, re-activating the corresponding conjunction neuron (**Fig.1B.5**), and therefore also the other features that were associated with it (**Fig.1B.6**). The object’s features are therefore recalled by auto-associative pattern completion, which brings them back into an attended, foreground state. Separate objects must always be encoded sequentially, which we suggest is plausible in light of the empirically observed attentional bottleneck in feature binding (Reynolds and Desimone, 1999).

To demonstrate the power of the model, we simulated a common visuospatial WM task (**Fig.2A**) in which participants remember the orientations of a set of colored bars (e.g. Gorgoraptis et al., 2011; Pertzov et al., 2016). Neurons were modelled as firing-rate units obeying a Hebbian plasticity rule (see **Methods**). Memory items were composed of combinations of features, and up to four unique items were presented sequentially to the feature units. After a delay, we probed one of the items by activating its color-feature alone, and recording whether its orientation was subsequently re-activated. Remarkably, just four color, orientation, location and conjunctive neurons each are needed to explain a wide range of behavioral and neurophysiological data, which no models have yet captured (**Table S1**).

**Fig.2:**
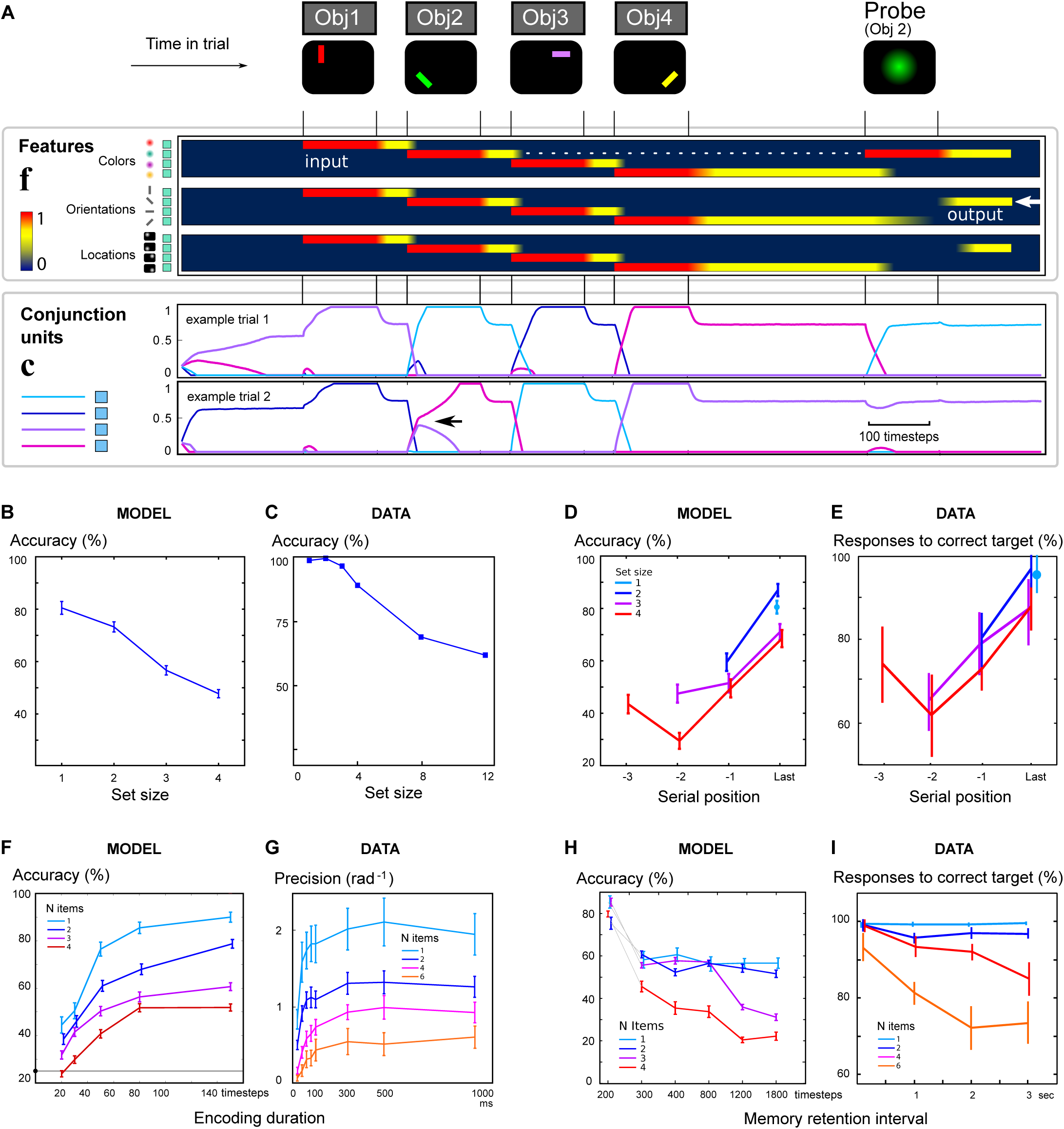
Predicting visuospatial WM capacity, encoding and decay. **A** To simulate WM performance, four objects are presented sequentially, by activating feature neurons (**f**, activity depicted as a heatmap from dark blue to red) indicating the color, orientation and location of each item. Conjunctive units (**c**) are shown below as four differently-colored traces. Conjunctive units compete to become active for each object. One conjunctive unit wins for each object, driving activity that persists even after input is removed (yellow parts of heatmap). At the time of the probe, a single feature is stimulated, triggering pattern completion. Recall is accurate if the orientation of the corresponding item is re-activated. Two example trials are shown; note that different patterns of conjunctive units are activated on different trials even for the same stimuli, depending on trial history. Example 1: good encoding. Example 2: weak encoding of the second item. Two conjunctive neurons with similar recent preferences compete to encode object 2 (arrowhead). When it is probed, item 4 is reported instead. **B & C** When more items are encoded in the model, recall accuracy is reduced, as observed in data (adapted from Luck and Vogel, 1997). **D & E** The last item encoded in the model is recalled better than others, as it remains active in the focus of attention during the delay period, matching observed serial order curves. Figure adapted from (Gorgoraptis et al., 2011) indicates the probability of reporting the target item as calculated by fitting the distribution of responses in a similar task. **F & G** Shorter encoding durations reduce modelled recall accuracy. Data from a similar task (adapted from Bays et al., 2011) where adding items reduced both initial encoding rate and asymptote. The model qualitatively reproduces the interaction observed in human performance. **H & I** The model predicts faster memory decay when more items are stored. This matches the empirical interaction between memory-set size and delay. Data adapted from (Pertzov et al., 2016) shows the modelled probability of reporting the target.

Crucially both the activation and learning equations were implemented continuously over a block of trials, with blank input in between trials, so that encoding, recall and interference from the previous trial all arose naturally from the way stimuli were presented. We tuned the model to perform at levels comparable to humans at this task (see **Methods**). For clarity, here we elected to keep the model’s operation almost identical for all the simulations, even though the experimental data we match come from a variety of tasks and measures.

### 2. Capacity limits and serial order in WM

A key feature of WM is its limited capacity. The more items held in memory, the less accurately they are remembered (Luck and Vogel, 1997; Bays and Husain, 2008). Simulated recall accuracy (**Fig.2B**) matched the set-size effect from classical visuospatial WM experiments (**Fig.2C**). This is because each additional stimulus competes for conjunctive neurons, and may corrupt or overwrite synaptic traces of previously-seen objects. Whether a previous item is overwritten is determined by the how well the currently-active features match the existing synaptic weights, which are themselves continuously subject to Hebbian rules. Therefore in our model, capacity is limited by interference between items in memory, in line with convergent evidence from multiple WM domains (Almeida et al., 2015; Farrell et al., 2016; Oberauer and Lewandowsky, 2014). Note that accuracy is lower than human data because the model chooses between four options rather than two, but varying the model parameters can make it arbitrarily more accurate (**Fig.S8,S9**). Importantly the model predicts the counterintuitive finding that storing extra features within a single object either occurs automatically(Allen et al., 2006) or else may incur no extra cost (Luck and Vogel, 1997). In fact our model predicts that in some situations a benefit can be observed for adding a new irrelevant but distinguishing feature to each object ( **Fig.S7**).

### 3. Serial order effects

When we remember a sequence of objects, we recall the first and last objects better (primacy and recency). Our model can reproduce both of these effects. Simulated performance ( **Fig.2D**) matched the serial position curve obtained in WM experiments (**Fig.2E**). The simulation suggests that neutrally, primacy benefits arise because the first object in a trial does not need to compete with ongoing persistent activity from a previous item (**Fig.1B4**). In our model this relies on the fact that, at the start of each trial, feature units are inhibited but previous synaptic weights are not erased – though there is no explicit signal to forget items from the previous trial. Recency benefits arose for two reasons: the finally-encoded item did not incur retroactive interference from subsequent items, and was already in an active rather than silent state at recall.

### 4. Encoding and maintenance

The time-course of encoding was interrogated by presenting items for brief durations, and demonstrated exponential saturation with an asymptote dependent on the number of items encoded. In a similar empirical study (Bays et al., 2011), memory precision (1/standard deviation of response angular error) followed a similar pattern. In that study, the probability of choosing the target was not calculated, but their reported precision appears to correspond well to our model’s probability of reporting the correct target orientation (**Fig. 2F&G**).

Simulations demonstrated that memory deteriorates faster when increasing numbers of items are remembered **(Fig.2H&I)**, as shown in a recent study (Pertzov et al., 2016). This arises because a greater proportion of items are held in an unattended state. Unattended items are more vulnerable to interference, because their synapses are gradually weakened over time according to the plasticity rule. Our model also makes the strong prediction that an item stored in an attended state (e.g. the final item in a sequence) is more robust to decay over time.

### 5. Shifting the Focus of Attention

An important advance over other models, is the ability of our model to re-activate a previous item by bringing it into the focus of attention. The logic here is that sensory input can guide attention by pattern-completion. In behavioral experiments, an “incidental” task inserted into the memory delay can shift attention to one of the items in memory (**Fig.3A**) (Zokaei et al., 2014a) bringing it into the foreground. We simulated “retro-cueing” one of the items during the memory delay by presenting one of its features for a brief period, which brought that item back into the focus of attention (**Fig. 3B**). The external cue could thus re-activate a memory item which was previously encoded silently. Note that this simulation illustrates how feature-selective units can exhibit task-dependent modulation because they also receive non-sensory input through rapidly-plastic synapses from the conjunctive units.

**Fig.3:**
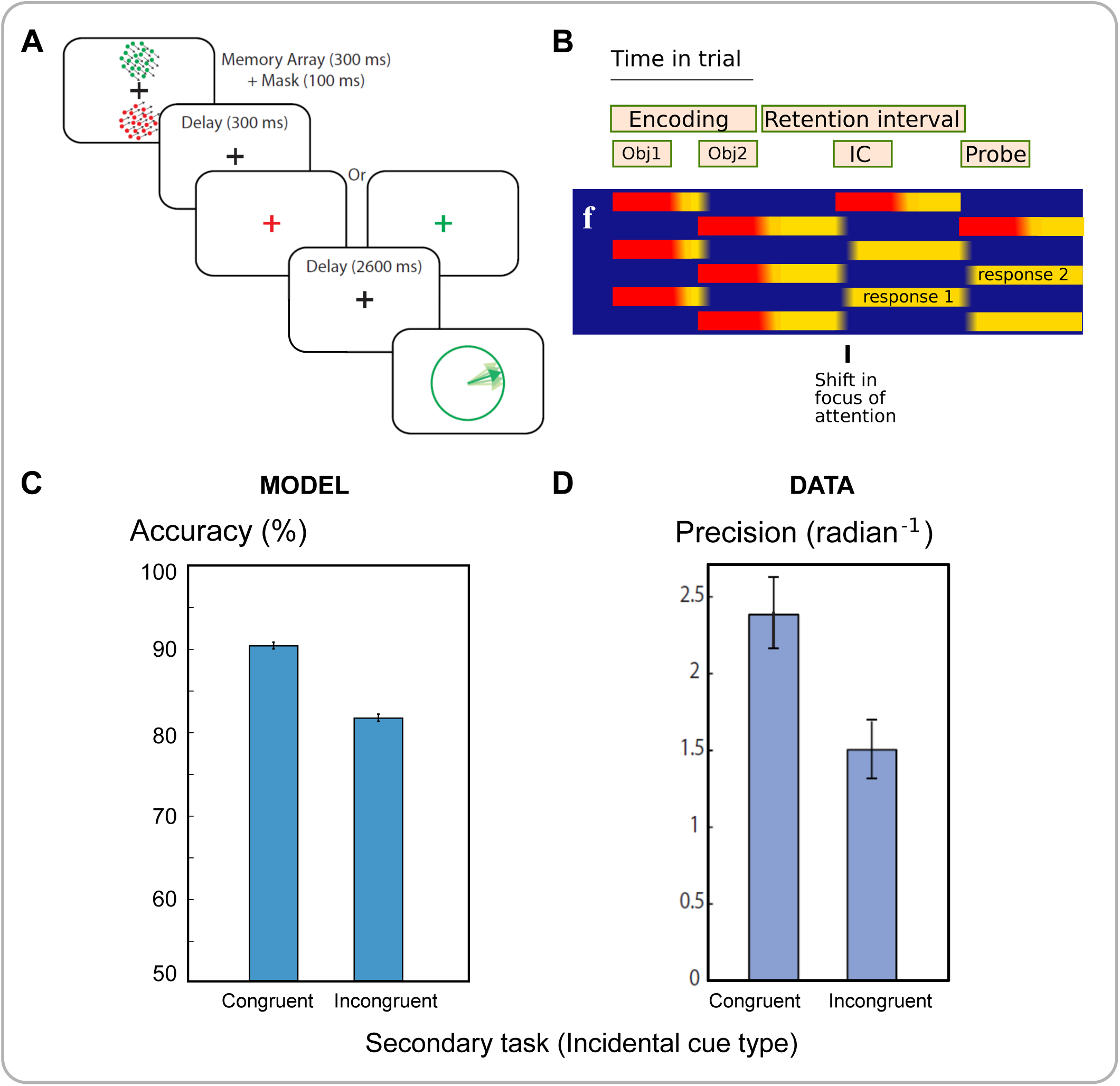
Shifting the focus of attention in WM. **A** Experiment (Zokaei et al., 2014a) where participants remembered two items, each comprising three features: color, location and orientation. During the retention interval, a color was shown, and as a secondary task, the location of the corresponding object had to be recalled. At the end of the delay, a color was shown which could indicate the same (“congruent”) or different (“incongruent”) object than the one tested during the delay. Participants then reported the orientation of the corresponding object. Reproduced under the terms of the Creative Commons Attribution 3.0 Unported (CC BY 3.0) license (https://creativecommons.org/licenses/by/3.0) from figure 1A of Zokaei et al. 2014a, The Journal of Neuroscience. January 1, 2014. 34(1); 158162. **B** Similar events were simulated, with an incidental cue (IC) during the delay. If the first object was cued, then persistent delay activity shifted to the cued item. **C&D** The model predicts that the item in the focus of attention before recall is reported more accurately, matching data. Reproduced under CC BY license from figure 1B of Zokaei et al. 2014a.

Recall of the incidentally-cued item improved, compared to the uncued item ( **Fig.3C**), matching experimental data (**Fig.3D**). This attentional shifting also explains how cues that indicate which item will be probed (predictive retro-cues, Rose et al., 2016) improve performance, even paradoxically when controlling for the retention interval’s duration (Myers et al., 2017).

### 6. Recall

After the probe feature was activated, it took a number of time steps for the conjunction and response feature units to become active. We measured this time to obtain reaction time predictions, which varied inversely with accuracy similar to empirical data (**Fig.S1**).

The process of recall may also be susceptible to interference, because it effectively uses pattern completion to re-activate the other features of the corresponding object. In particular, the memory probe itself can interfere with recall, for example if it contains a feature on the dimension that needs to be reported (**Fig.S2**), in line with empirical probe-interference effects (Souza et al., 2016). Interference of another kind arises when recalling items as a whole series: often the preceding or following item is reported instead (Smyth, 1996; Solway et al., 2012). Although our simulations probe a single item at a time, they still demonstrate such “transposition errors”, where consecutively presented objects are confused (**Fig.S3**).

### 7. Neural encoding of items in WM

Three major predictions emerge about neural decoding. First, an emergent property of our framework is that sustained activity represents a single item held in memory (Funahashi, 2017), but not multiple items (Lara and Wallis, 2014). We used a linear decoder to extract information about one feature of one of the items in WM, after items had been encoded. The predictions of the model for decodability from feature-selective neurons (**Fig.S4**) are in keeping with human and nonhuman physiological data demonstrating that only the attended WM item is decodable using standard techniques (Konecky et al., 2017; Lewis-Peacock et al., 2012; Sprague et al., 2016). Second, evoking neural activity by stimulation can restore decodability from EEG signals (Rose et al., 2016; Wolff et al., 2017). We simulated transcranial magnetic stimulation (TMS) by an indiscriminate pulse of activation to feature neurons ( **Fig.4A)**, and decoded one feature dimension from feature-selective units (**Fig.4B**). If the model’s color and orientation feature dimensions are considered as mapping to spatial location and stimulus category respectively, then the simulation matches the effects of TMS on decoding (**Fig.4C**) (Rose et al., 2016), or if they are instead mapped to spatial location and orientation, then the model’s results reproduces the effects of a high-energy visual pulse (Wolff et al., 2017). Simulating a stronger pulse of stimulation disrupted attention, but not synapses. This worsened recall of the attended item, yet contrarily improved unattended items (**Fig.4D&E**), precisely as demonstrated empirically (Zokaei et al., 2014a).

**Fig.4:**
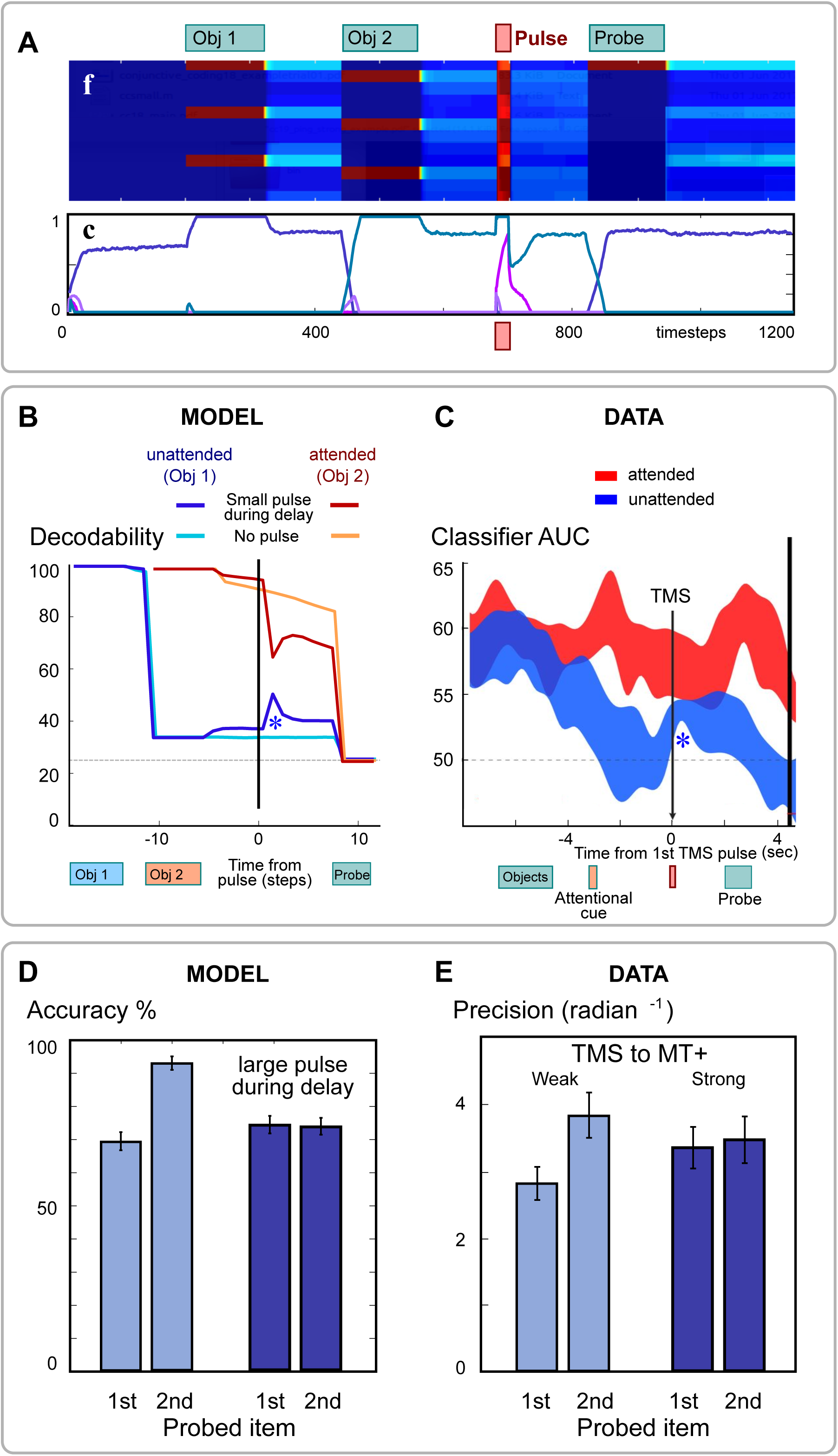
Introducing a pulse of excitation during the delay period. **A** After presenting two items, during the delay all feature neurons **f** received an excitatory input pulse **i**=+1, consequently activating conjunction neurons. **B&C** We tried decoding the identity of each of the two stimuli from feature neuron activity. Although the first object was not decodable without the pulse, it became transiently distinguishable (*) after the pulse. This matches the observed increase in decodability after TMS (Rose et al., 2016). **D&E** Stronger pulses altered model performance, abolishing the benefit for the second item, which was in the focus of attention. The pulse disrupted persistent activity, re-instating competition between conjunctive neurons. The prediction matches observed effects of TMS targeting motion-selective cortex (Zokaei et al., 2014a).

Third, the model predicts that decoding from prefrontal cortex is unreliable (Lee and Baker, 2016). This is because the concept of a receptive field breaks down for conjunctive neurons. The same activity can have *different meanings* on different trials, dependent on residual synaptic weights from previous trials. Such neurons should show much stronger representations over short timescales. We predict this will manifest behaviorally, with better recall for a feature combination present on the previous trial (**Fig.S5**), because the same conjunction unit will be reused. Moreover, neural activity patterns in conjunction neurons predict stimuli strongly if we consider data only from *contiguous* pairs of trials, compared to data from temporally-separated trials (**Fig.5A**), and the pattern similarity should be even lower when intervening stimuli involve a recombination of the features (**Fig.5B-D**). This confirms that each conjunctive neuron’s activity represents different things, as its synaptic weights change. Such a system can flexibly encode a broad variety of novel information rapidly, without incurring the combinatorial explosion that haunts previous fixed-selectivity models (Matthey et al., 2015; Postle et al., 2006).

**Fig.5:**
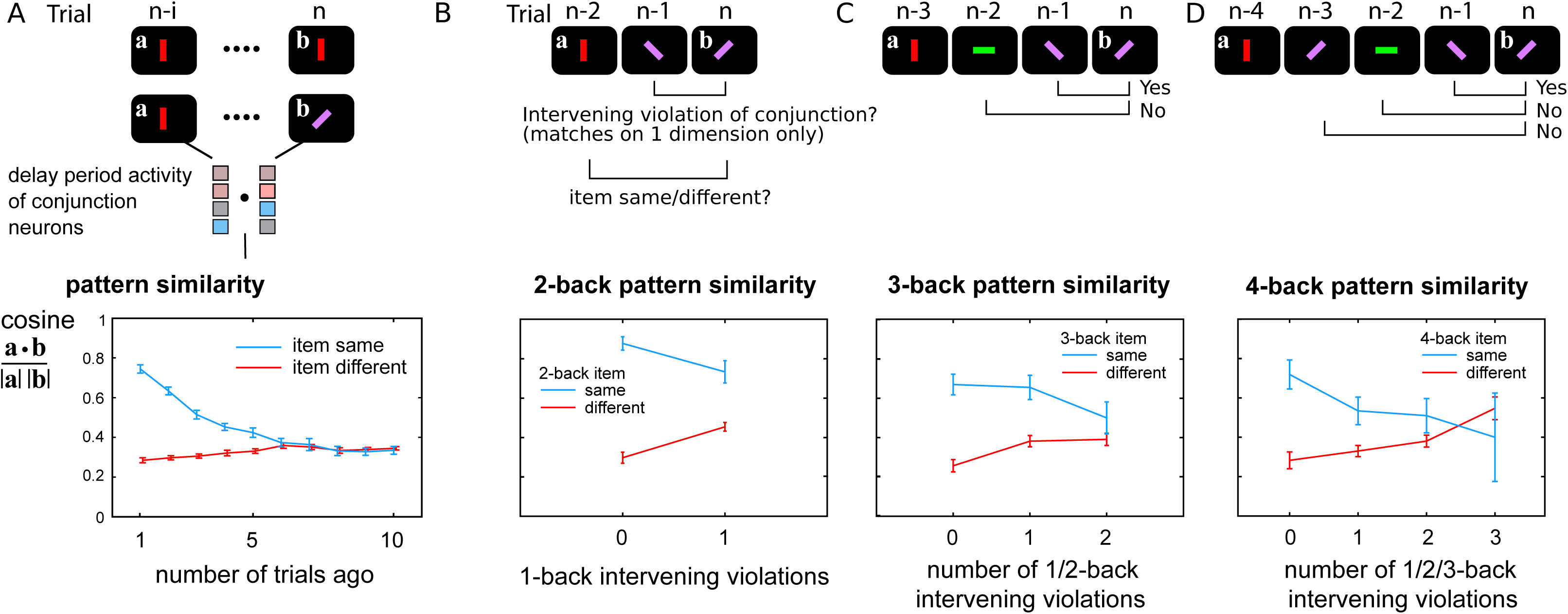
Conjunctive unit representations are stable over short timescales. Conjunctive units change their selectivity over short periods. If selectivity were stable, neural patterns should be similar when the stimulus is the same. We compared similarity of the pattern of an earlier trial, to trial n, during the delay periods of a series of 1-item trials. **A**) The similarity of the conjunctive neurons’ delay activity pattern is calculated for trials where the stimuli were identical (blue line) or different (red line). Patterns were more similar when stimuli were the same, compared to when stimuli were different, indicating “classical encoding” at least for nearby trials. This classical behavior decreased with the temporal distance between trials. Since we modelled the extreme case where neurons are *purely* conjunctive, with no feature selectivity, consistency of pattern is completely abolished after about 6 trials. **B-D**) The model predicts that interference reduces pattern similarity over time by overwriting the synaptic weights. If the objects in intervening trials share one feature with the nth trial object, but mismatch on the other feature dimension, then we say the conjunction between the two feature dimensions is “violated”. **B**) When the intervening trial contained a violation, the patterns on the n-2 and nth trials reflected the stimuli much more weakly, indicating interference or overwriting of the original conjunction. **C** and **D**) Trials 3-back and 4-back were similarly examined, this time asking how many intervening conjunction violations occurred. The more overwriting that occurred between the n-3 and nth trials, the less classical encoding could be observed.

### 8. Simulation of task sets

The same system can also implement stimulus-response rules, if some feature neurons represent motor plans. In this case, we encode a *task rule* by attending to a stimulus and a motor plan together. For example, if a left-hand movement plan is activated while a red color-feature is simultaneously activated, they will be encoded together into working memory. The conjunction of sensory features with a motor plan creates a task-set mapping (Duncan et al., 2012). Later, that stimulus can also re-activate the corresponding motor plan by pattern-completion, triggering the movement – so that the stimulus generates a response. Task sets can therefore be rapidly formed by sequentially attending to stimulus-response pairs (Curtis and D’Esposito, 2003), and deciding on an action is simply the motor analogue of WM recall.

To simulate stimulus-response mapping, we presented the task rules sequentially, each consisting of a pairing between one color and one response (**Fig.S6A**). Then on each subsequent trial, a single color from the set was shown, and the response was recorded. The model reproduces Hick’s law, in which response times are longer in situations when more response options are possible in the current task set (**Fig.S6B**) (Proctor and Schneider, 2017). It also produces faster reaction times when the response is repeated from the previous trial ( **Fig.S6C**), in line with experimental evidence (Schvaneveldt and Chase, 1969).

In this situation, the role of prefrontal conjunctions can be viewed as *controlling* representations in posterior cortex, i.e. routing information from perceptual to motor representation as governed by task sets held in working memory, a role classically assigned to executive/supervisory attention. Critically the model predicts that, because the task rules are held in WM across many trials rather than being repeatedly overwritten, the current stimulus and response (i.e. the active task rule) are consistently decodable from conjunctive neurons, until the rules change ( **Fig.S6D**). This contrasts with WM storage, where frequent overwriting leads to poor decoding, and may explain why task rules have generally been easier to decode from PFC (Reverberi et al., 2012; Sakai, 2008).

## Discussion

The model of freely-conjunctive neurons presented here accounts for both sustained firing and activity-silent synaptic traces in WM (Silvanto, 2017; Stokes, 2015), and consequently makes a range of testable behavioral and neural predictions (**Table S1**). This neuronal framework provides a parsimonious mechanism for feature binding, general-purpose memory ‘slots’, and task sets. The model reproduces classical WM effects of capacity, serial order, encoding rate, temporal decay (**Fig.2**), reaction times, and transposition errors (Fig.S1&3), as well as the ability to switch attention between items within memory – a phenomenon that evades most current models (**Fig.3**). At a neural level, it explains why it is difficult to decode memory contents from prefrontal activity, why only the item in the focus of attention can be decoded elsewhere (**Fig.S4**). Further it explains why decodability can be restored by re-focusing an unattended item, or after a perturbation such as transcranial magnetic stimulation (TMS) or bottom-up input (Rose et al., 2016; Wolff et al., 2017), which presumably re-activate the conjunctive neurons and thus an object’s features through synaptic traces (**Fig.4**). The model also makes strong novel predictions about probe interference, trial-to-trial effects (Fig.S2&5), and disruption of neural pattern similarity by intervening stimuli (**Fig.5**).

### Relation to previous models

Rapid plasticity has long been demonstrated in cortical neuron receptive fields (Edeline et al., 1993) and may arise through a variety of synaptic mechanisms (Zucker, 1989; Tsodyks and Markram, 1997; Fischer et al., 1998; Dittman et al., 2000; Jensen et al., 1996; Malsburg, 1981). It has previously been suggested to underlie short-term retention of information in a network of selective and nonselective neurons (Mongillo et al., 2008), similar to theories of hippocampal binding in long-term memory (Burgess and Hitch, 2005; Rizzuto and Kahana, 2001). Rapidly plastic networks account well for serial recall (Farrell and Lewandowsky, 2002; Fiebig and Lansner, 2017; Sandberg et al., 2003) but cannot distinguish between focused and unfocused items in memory, and do not explain sustained delay-period firing (Funahashi, 2017). Sustained activity models may also account for some attentional effects on decodability (Schneegans and Bays, 2017) but cannot reinstate information that becomes fully undecodable.

To support flexible attractor states, we postulated two distinct modes of neural representation (**Fig.1**). First, feature-selective neurons are traditional, place-coded (“labelled-line”) units. They are selective because they have some fixed, non-plastic inputs (or in the case of motor units, fixed outputs). But if plasticity modifies both the input and output synapses of a neuron, the meaning or interpretation of a neuron’s firing will also change. This is simply because neurons *code* information only in virtue of their inputs and outputs. Plasticity therefore begets a new category of flexibly-coding neurons, where the information signaled by firing is protean and dependent on the history on each trial. Decoding the fine-grained identity of stimuli from prefrontal cortex is unreliable compared to posterior sensorimotor regions (Cogan et al., 2017; Lee and Baker, 2016), because the idea of a receptive field breaks down. Standard decoding methods assume trial-to-trial stability of activation patterns to represent a given feature, and so do not measure the sequential effects we predict. This flexible coding scheme is crucial for our model to generate two phenomena. First, it permits sustained activity that is guided dynamically by task sets or objects in memory, which we postulate corresponds to attentional interactions between frontal and temporo-parietal regions. Second, because individual neurons can encode different things at different times, information must *compete* to be encoded by any conjunctive neuron – thus leading to a capacity limit for general-purpose information storage, observed in both WM and attention. This may help resolve a long-standing theoretical debate on whether working memory consists of pointers, or activated long-term memory (Norris, 2017): conjunctive neurons act as pointers that activate long-term memories.

### Relaxing the model’s assumptions

In this study we deliberately chose to study the simplest possible model that could support conjunctive neurons. The very small number of neurons, and their simple learning and dynamics, makes it much easier to see how they interact to generate the novel predictions. Moreover it is much more transparent where the model can or cannot match existing data. Naturally there are many directions in which the model needs to be extended, to fully reproduce the phenomena observed in real neurons. A number of its assumptions can plausibly be relaxed.

#### 1. Pure flexible and stable representations

For simplicity we have treated conjunctive neurons as “pure”: i.e. that they are homogeneous and domain-general, resulting in inability to decode information across many trials. This is certainly implausible because all-to-all connections between PFC and feature-selective neurons are not feasible. Moreover, how can we then explain studies that *do* demonstrate decoding of WM from prefrontal areas? In reality, we envisage that each conjunctive neuron is likely to receive inputs from only a subset of feature neurons. In order for conjunctive neurons to bind features into objects, these inputs must at least include multiple feature dimensions *and* multiple features in each dimension. The model is therefore potentially compatible with the presence of mixed selectivity (Rigotti et al., 2013), which would provide a background of weak input selectivity based on the presence or absence of connections, upon which rapid plasticity is superimposed.

Further, there may also be significant topography in conjunctive cells connectivity. For example, different regions of prefrontal cortex may be specialized for remembering different kinds of information (Romanski, 2004). This may have two desirable consequences. First, aspects of the attended object – especially information that is highly topographical in posterior areas, such as stimulus category and spatial location – would be consistently decodable from prefrontal cortex (Lee and Baker, 2016) but will be modulated by relevance (Kornblith and Tsao, 2017). Second, conjunctive neurons in different prefrontal subregions may connect preferentially to visual, motor or auditory cortex, which could account for the separability of visuospatial and phonological WM and also their overlap (Morey et al., 2011). We note that stable mixed selectivity, even without plasticity, could in some situations produce binding and capacity limits (Matthey et al., 2015). However without additional mechanisms, it would presumably not account for attentional shifts, activity-silent storage, or apparent control over posterior cortical areas, and moreover makes it challenging to internally ‘read-out’ WM contents.

We treated “features” as just simple perceptual attributes, but we believe that our class of feature-selective neurons could include any aspect of the world that is encoded in a stable way, including those aspects that incorporate long-term knowledge, such as object identity, category, or even linguistic information such as word meanings. These attributes are likely to be encoded stably in posterior cortical areas, in contrast to the temporary combinations of information represented in an ephemeral way – e.g. for online manipulation – as typified by our conjunctive neurons. The current simulations used only a single, rapid learning rate, but it remains to be studied how this could be reconciled with longer-term learning.

#### 2. Internal control over attentional shifts

We have assumed that attentional shifts are externally cued. Endogenous shifts of attention are not modelled. One way of implementing internally-generated attentional modulation would be to de-stabilize the persistent activity by adding delayed suppression, or refractoriness, to the competitive conjunctive neurons. The result would be that, after an object is attended, its activity is extinguished after a delay, leading to a transient and unstable focus of attention. Akin to some models of visual attention guidance (Itti and Koch, 2001), attention will then be successively redeployed towards the weakest-represented features in WM. This could potentially account for four key phenomena: (a) rehearsal, in which attention moves sequentially between items during a memory delay, (b) the ability to free-recall WM items in order, (c) the guidance of visual search, and (d) in our model, to permit serial encoding of a simultaneously-presented memory array.

Although WM maintenance commonly engages PFC, evidence from neuropsychology and functional imaging suggests PFC’s role includes cognitive control, WM manipulation, and response selection, rather than simply WM storage (Bechara et al., 1998; D’Esposito and Postle, 1999; Rowe et al., 2000; Thompson-Schill et al., 2002), and it remains to be tested whether the conjunctive neurons we propose can perform such functions. For example, we cannot account for the ability to “gate out” distractors, and prevent them from being encoded in WM. If sensory input is sufficiently weak, in our model, it can cause transient activation of conjunctive neurons without capturing attention, such that the ongoing attractor state remains stable. But how could *irrelevant* distractors be ignored, while still allowing relevant inputs to capture attention? To achieve this, conjunctive units would themselves need to be under higher-level control. The current model, with only one layer of conjunction units, does not explain higher order control of attention, since sufficiently-strong bottom-up stimuli that match a conjunction will always tend to re-activate that conjunction and thus capture the focus attention. The conjunction and feature neurons together simply act as a “matched filter” amplifying patterns that have recently been active (Chrysikou et al., 2014; Hayden and Gallant, 2013). Perhaps gating vs granting access to working memory by preventing this might be controlled by interactions between prefrontal cortex and the basal ganglia (Badre, 2012; Chatham et al., 2014).

#### 3. Location of conjunctive neurons

Conjunctive-coding neurons might not be confined to prefrontal cortex. Other regions that play a role in working memory, such as the hippocampus, basal ganglia or thalamus, might also contain freely conjunctive neurons. Moreover, there may be a continuum or overlap of mechanisms subserving working memory and episodic memory (Fiebig and Lansner, 2014). However the volatile synaptic weights we propose would produce strong but evanescent trial-by-trial selectivity changes (**Fig.5**), quite unlike the rapid but long-lasting associations proposed in the hippocampus. A more intriguing possibility is that both freely-conjunctive and stable-feature neurons are actually present in the *same* brain regions, with a spectrum between highly-plastic and stably-coding neurons.

#### 4. Spiking and synchrony

Neurophysiological evidence points to synchrony of neuronal firing as a key feature of attention (Myers et al., 2017). Since we only simulated firing rates, synchrony and oscillations are not observable. Our reciprocal activation and “reverberatory” sustained firing could lead to spiking synchrony that is cross-region and attention-dependent. The current model makes no specific predictions regarding theta- and gamma-band activity. To generate such predictions, a spiking model, possibly including the thalamus and intracortical oscillations, may be needed. This is far from trivial and further study is required to determine if a spiking implementation of this network would generate the same predictions. An open question is how this might be reconciled with evidence that multiple items may be stored in an electrically active state. For example, could several attractors be simultaneously active, kept distinct through inhibition (Wei et al., 2012) or multiplexed by phase relative to oscillations (Jensen et al., 2002; Lisman and Jensen, 2013) ? Our model is incompatible with these possibilities.

In summary, a single architecture captures both persistent activity attractors and silent synaptic memory. We introduce a new scheme of transient flexible neuronal coding, that can support many empirical phenomena (**Tables S1/2**) including the “focus of attention”, and generates numerous testable neural predictions.

## Acknowledgments

Research was funded by an MRC Clinician Scientist Fellowship to SGM, a Wellcome Trust Principal Fellowship to MH and NIHR Oxford Biomedical Research Centre. SGM conceived the model, ran simulations, and wrote the manuscript. TV advised on modelling, NZ, SJF and MH co-wrote the manuscript. We thank Prof. Mark Stokes and Dr. Nicholas Myers for comments on a draft, and Dr Eva Feredoes whose discussions inspired some of the concepts.

## Supplementary Materials

### Methods

The present model considers a minimal arrangement for three feature dimensions, each with four possible feature values, allowing 12 features to be encoded. Each feature unit receives input when a particular feature is present in the stimulus. Four conjunctive units are fully connected reciprocally to the 12 feature units. These connections are all excitatory, and initialized to be random. Four conjunctive neurons is the minimum possible number that can bind information from four objects. The fully-connected network therefore required 48 weights to conjunctions from features (**W**^*cf*^) and 48 to features from conjunctions (**W**^*fc*^). The activity of all units **f** and **c** were initialized to zero at the start of simulation, and weights **W**^*fc*^ and **W**^*cf*^ were randomly assigned from a uniform distribution over the interval [0, 1]. In its simplest form, the activity update equation was:

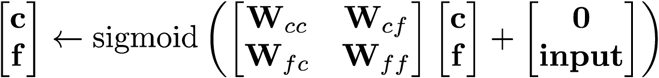

The conjunctive-feature synapses **W**^*cf*^ and **W**^*fc*^ were updated by a Hebbian covariance rule, whereas the fixed inter-conjunction (**W**^*cc*^) and inter-feature (**W**^*ff*^) synapses each comprised two components:

1. blanket lateral inhibition between conjunction neurons or between features within the same dimension, to implement competition, and
2. self-excitation, so that firing does not stop suddenly when external input is removed, but rather decays exponentially with time.

So the update equations can be written out more fully as:

neuron activation:

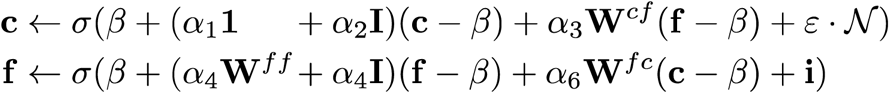

synaptic updates

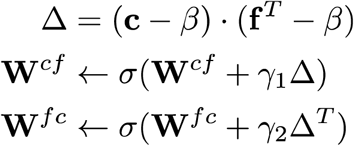

constants:

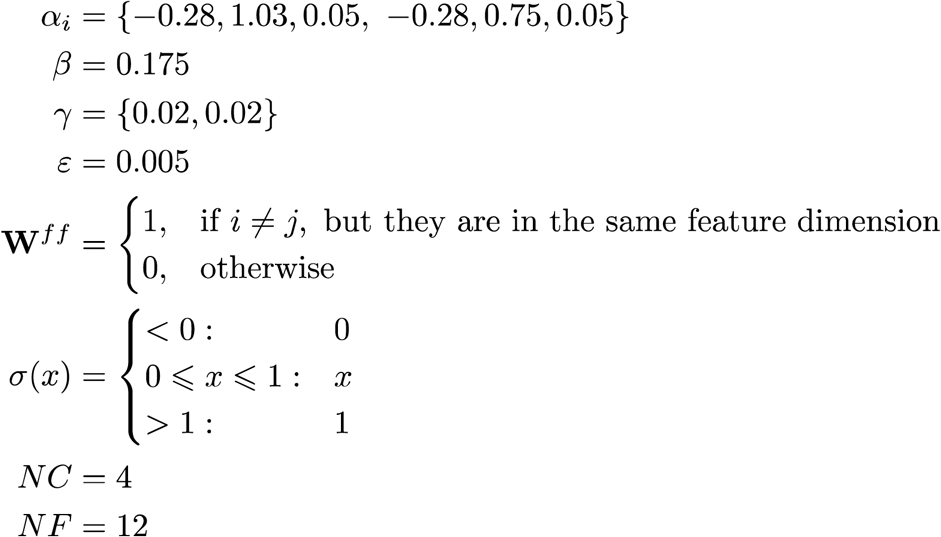

In these equations, **c** and **f** are the activities of conjunctive neurons and feature neurons, and **i** is the external input. The six synaptic free parameters α were:

α1, α4: mutual lateral inhibition between neurons
α2, α5: self-excitation or temporal decay, and
α3, α6: synaptic gain for the conjunction-to-feature and feature-to-conjunction synapses,

for the conjunctive and feature neurons respectively. β is the baseline neuron activity, *γ_i_* are the learning rates, and ε is the amount of noise added to the conjunctive units. The function σ constrains values lie between 0 and 1 and, for simplicity, was chosen as: σ(y)=min(1, max(0, *y)).* The identity matrix **I** produces self-excitatory synapses, and a matrix of ones (**1**) produces lateral inhibition. In simulations, the 12 features were arranged into 3 dimensions (color, orientation and location), so that the feature-to-feature inhibition (**W**^*ff*^) was arranged in 3 blocks of 4 units. *N* indicates a gaussian white noise vector with s.d.=1. We constrained learning rates to be identical in both directions (*γ*1=*γ*2), and noise was in fact not required in order to obtain the typical patterns of errors reported here (ε=0), giving effectively 8 free parameters. A minimal algorithm to reproduce the main result is provided at the end of this section.

#### Implementation of stimuli and simulations

Model equations were simulated in MATLAB (code available at [www.smanohar.com/wp/wm/download.html]). A web-based interactive simulation can be accessed at http://www.smanohar.com/wp/wm. Each run simulated 200 trials. To test different hypotheses, the simulation setup was varied from a canonical setup. The canonical simulation resembled common working memory experiments (Bays and Husain, 2008). A series of objects was presented, followed by a memory probe that triggers recall. Each object excited one feature in each dimension (colour, orientation and location). The features of the objects presented on each trial were random, with the constraint that no two objects within one trial shared any features. Each trial started with a foreperiod for equilibration lasting 200 time steps, with the features inhibited (input **i** = -1). Then during the encoding epoch, each memory item was presented by activating the corresponding feature units, by maximally activating the features present in the object (i = +1) and inactivating the absent features (i = -1). Stimuli were presented for 120 time steps, followed by an inter-stimulus interval of 50 time steps with no input ( **i**=0). After the last item was presented, the memory delay period of 240 time steps followed, with no input, mimicking a retention interval. Then at the probe epoch, the feature acting as the retrieval cue was activated (i=+1), and the other features were inhibited (i = −1). The probe lasted 120 time steps. Finally, the response interval constituted 220 time steps with no input. This permitted re-activation of the features to be recalled. The feature that reached the highest level of activity during this period was selected as the response. In the case of an exact tie-breaker the decision was randomized. An example of this sequence of events shown in **Video S3**.

Parameter selection was performed initially by trial and error to achieve the hypothesized dynamics. First, the decay, inhibition and input weightings α, for the conjunctive and feature neurons were adjusted to create a sustained activity plateau and to ensure that conjunctive neurons competed in a winner-takes-all manner. Then the learning rates *γ*, were adjusted to allow sufficiently rapid weight changes such that over 50 time steps, a trace remained that represented which combination of feature units had been active. The baseline was approximately 20% maximal to permit deactivation of neurons, so that conjunction units that lose the competition will ‘unlearn’ their associations (Stanton and Sejnowski, 1989). A small amount of noise was added to ensure conjunction units were never identical, to facilitate symmetry-breaking.

Certain effects were more or less sensitive to changes in model parameters. Overall accuracy could be made to range from consistent 100% to consistent 0%, depending on choices of α, β, γ, and ε. We aimed for 70% overall accuracy, allowing a wide dynamic range of performance to examine the predicted effects, so this was the regime in which the main simulations were run.

#### Simulation 1: set size

Simulation parameters were as above, with *α_i_* = { -0.28, 1.03, 0.05, -0.28, 0.75, 0.05 }; β=0.175, γ = {0.02, 0.02}; ε = 0.005. Timings were: foreperiod=200 steps, objects=120 steps, inter-stimulus=50 steps, retention delay=240 steps, probe=120 steps, recall=240 steps. Trials proceeded as above, with four items presented sequentially. One, two, three and four items were presented in different trials, and each serial position was probed. For multi-item sequences, each item in the sequence was probed equally often. This gave ten (1+2+3+4) trial types, and 200 trials of each type were simulated. The average accuracy over all serial positions was calculated for each set size (**Fig.2B**). Error bars are the standard error of the accuracy when trials were broken into subsets of 20 trials each. Simulations were performed both using interleaved trials, and also with one condition per block, and comparable results were obtained with both methods.

Capacity limits were naturally limited to four in this simulation because only four conjunctive neurons were present. However the capacity limit does not *directly* relate to the number of conjunctive neurons, but rather, to the *proportion* of conjunctive neurons that will become simultaneously active during winner-takes-all competition (¼ in this case). This is in turn determined by the level of inhibition. A simulation using 8 conjunctive neurons also reproduced the set-size effects, with inhibition tuned to allow two neurons to be active at once.

Note that absolute modelled accuracy was lower than in the empirical data, because participants made same/different judgements (chance=50%) whereas our simulations recalled actual features (chance=25%).

#### Simulation 2: serial position

The run was identical to simulation 1 above. Trials were grouped according to the serial position of the probed item, and by set size (**Fig.2D**). Accuracy was calculated as per simulation 1. Note that in all simulations, feature unit activity was held at zero at the start of each trial ( **i** = −1) to simulate a foreperiod or inter-trial interval. This permitted the first item of the sequence to be encoded faster. We then explored different parameter sets, varying α, β, γ and ε. The recency effect was robust across a wide range of parameters, and was often strong enough to push performance to 100% for the last item. The presence of a primacy effect was more strongly dependent on the specific presentation timings and on the choice of α_*i*_. This matches empirical studies of visual WM, which do not always find primacy effects.

#### Simulation 3 – encoding duration

The time that each item was presented for was varied. This ranged from 10 to 120 time steps (in increments of 20 ms), with all items being presented for the same duration. The inter-object duration was fixed at 50 ms as in previous simulations. The average accuracy was collapsed across all serial positions for each set size, and plotted as a function of encoding duration (**Fig.2F)**. The simulation demonstrates an interaction between set size and time, such that when more items are stored, the decay is faster. In order to avoid floor effects for this simulation, where accuracy in the 4-item condition could approach chance very quickly as the encoding duration decreases, we increased the ‘stickiness’ of conjunction units in the model, by reducing the decay factor α1 and uptitrating their input gain from each other (α_2_) and from the feature units (α_3_). We therefore set

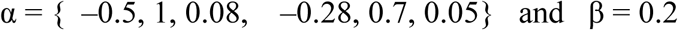

for this simulation, and the consequence was that the overall accuracy of the model increased from 75% to around 90% while preserving the set size and serial position effects. This ‘higher performance’ regime was used for simulations 3, 4 and 5. Other than improved overall accuracy, these parameters produced qualitatively similar effects to the primary simulations (**Fig.S9**).

#### Simulation 4: effect of memory delay

We varied the number of time steps after the final item was presented, until the onset of the probe. The delay varied between 200 and 1800 steps. For each duration, simulations of 200 trials were run for 1, 2, 3 and 4 items, with each possible position being probed (i.e. total 2,000 trials per duration). In order to avoid floor effects, where accuracy in the 4-item condition could approach chance as the delay increases, we used the same regime as Simulation 3. Data is plotted as a function of the delay (**Fig 2H**).

#### Simulation 5: transposition errors

Here we studied the tendency to incorrectly report features from items temporally adjacent to the probed item. This arises because occasionally, the same conjunctive unit is activated for two consecutive objects, when the second object fails to sufficiently drive a different conjunction unit. In this case, features of two consecutive objects will be confused.

Data were taken from the standard 4-item condition where four items were presented and one is probed, using parameters of Simulation 3. For this simulation, trials were grouped according to serial position of the probed item. Four responses were possible on each trial, and for each trial, we take the serial position at which the reported feature *actually* appeared on that trial (**Fig.S3B**). This figure shows a histogram of the model’s responses. Over 2000 trials, the probability of making each of the four responses was calculated. Each line shows the probability of reporting the orientation of the four items presented in the sequence (x-axis), when a particular serial position was probed (each as a different line). We show logarithms of the mean error rate as in (Farrell and Lewandowsky, 2004), and added a small offset of 10^−3^ since some runs had zero errors. Error bars are standard error of the logarithm of mean error rate. The four possible responses are aligned such that the correct response appears at position zero, at the center of the graph (i.e. the *probed* item’s orientation). Sometimes the model erroneously reports an item previous to the one probed (x<0) or an item later than the item probed (x>0). The mean of the four probe conditions is shown in red.

#### Simulation 6: Incidental retrocue

In the empirical study, the primary task involved recalling the orientation of the item with a given color, and the secondary task (“incidental cueing”) required participants to report the location of the item with a given color, which could be congruent or incongruent to the ultimately-probed item. We simulated the empirical task (**Fig.3D**) by presenting two items, as per the 2-item condition in simulation 3. After the items were presented, a 120 time-steps retention period followed, then the probe feature (incidental cue) for one of the two items was activated for 40 time steps (“IC” in **Fig. 3E**). After a further 120 time steps corresponding to recall of the third dimension for this item, a further memory delay of 120 time-steps was included. Then the final memory probe was activated, and recall of the second dimension was measured as previously. This final probe could either be the same item (congruent), or the other item (incongruent), as the one cued for the first response. 200 trials were simulated for each condition. Accuracy was plotted for the valid and invalid incidental cue conditions ( **Fig. 3F**).

#### Simulation 7: Reaction time (RT)

RT was calculated by finding the time at which the winning feature reached its maximal value in the period after the probe. The time to reach 98% of maximum was used, rather than using an absolute threshold, because the final stable value of an activated feature differed when different model parameters were used. Using an arbitrary fixed threshold or the rate of rise yielded qualitatively similar but less consistent RT effects. To examine basic set size and serial position effects, the same trials were used as in simulation 1. Mean RT with standard error is shown (**Fig.S1A**), in comparison to data (McElree and Dosher, 1989) (**Fig.S1B**). RT was generally inversely related to accuracy.

#### Simulation 8: probe interference

Many working memory tasks have asked participants to adjust features of the probe to match the remembered features (Zhang and Luck, 2008). In these experiments, the probe contains a feature that is irrelevant to recalling the item. For example, if participants must report the orientation of a bar with a given color, then using a colored bar as a probe introduces an orientation feature that conflicts with the remembered item. The model predicts this will interfere with re-focusing the item (**Fig.S2**). To simulate this, sequences of 1 to 3 items were presented to the model, using an identical setup to simulation 1. However at the time of probe, two features were activated; one in the probe dimension, and one in the recall dimension. As previously, the probe feature input was +1, and other features on that dimension were -1. However an additional input +1 was added for one orientation feature. The additional feature, in the recall dimension, was one that had *not* been presented on that trial. Since only 4 features were present in this model, this constraint meant that we could only test set sizes of one to three items.

Performance was worse when the probe contained an interfering item. There is some empirical support that this might indeed be the case (Souza et al., 2016). The simulation predicts interference across all set sizes. Conversely, we can also predict an improvement in performance for probes containing an additional *helpful* feature, for example if participants must report an object’s orientation given both its color *and* its location.

#### Simulation 9: Weak TMS pulse reactivates delay period decoding

Here we enquired whether the model could reproduce the phenomena where unattended items, which are not normally decodable from brain activity, could be brought back into a temporarily decodable state by applying a pulse of activation. Empirically this has been demonstrated using a nonspecific high-energy visual stimulus pattern (Wolff et al., 2017), or by applying a TMS pulse to sensory cortex (Rose et al., 2016). In the simulation, two items were presented sequentially to the model, exactly as in simulation 1. During the delay period, the feature neurons received a flat high-valued input (**i**=+1) for 10 time steps. This was compared to an identical condition with no stimulus. The delay period therefore comprised wither 120 timesteps + 10 timestep pulse + 120 time steps, or in the no-stimulus condition, 250 timesteps (**Fig 4A**).

2000 trials were simulated, and half the trials were used to construct a linear classifier that predicts the identity of each of the two items presented on that trial. The two classifiers were tested on the remaining trials, to give the decoding accuracy. Decoding was performed across trials at each time point independently, indicating the degree to which neural activity in the feature units predicted the identity of each item. The first object presented was termed the ‘unattended’ object, since during the delay, attention was focused on the second (final) object. Decodability of the unattended item was transiently restored after the pulse ( **Fig.4B** dark blue trace), reproducing the phenomenon in the TMS study (**Fig.4C**) and using a high-energy neutral visual stimulus.

Two further predictions are that 1) stimulation of prefrontal regions should have similar effects, and 2) selectively stimulating specific feature neurons e.g. in sensory cortex will have stronger effects when those features are part of an item currently in memory – i.e. when the synapses from that feature neuron to the conjunctive neurons are already strong.

#### Simulation 10: Strong TMS pulse disrupts focus of attention

A strong TMS pulse to sensory cortex (MT+) has been shown to disrupt the focus of attention. To reproduce this, we used precisely the same simulation as in 9, but increased the pulse duration to 20 timesteps duration. We compared accuracy for probing the first item, vs the second item that was presented, with and without pulse. The presence of the pulse reduced accuracy for the second item, but paradoxically improved memory for the first item (**Fig.4D**). This is because the pulse indiscriminately re-activated the feature and conjunction neurons, disrupting the focus of attention but not the underlying memory traces. Indeed after the pulse, the attractor state sometimes shifted back to the first item. Precisely this phenomenon was observed in a TMS study (Zokaei et al., 2014a) (**Fig.4E**). The model predicts the same effects if prefrontal neurons are stimulated.

#### Simulation 11: Delay-period decoding

Three items were presented in sequence to the model. This was identical to simulation 1 except that the delay period duration between items was increased to 100 time steps, to test decoding during the stable attractor state. Each serial position was probed on 2000 trials. As in simulations 9 and 10 above, half the trials were used to construct linear classifiers that predict the identity of the item that was shown at each given serial position on a trial. The three classifiers were then tested on the remaining trials, to give the decoding accuracy. Decoding at each moment in a trial indicates the degree to which neural activity in the feature units predicted the identity of the item presented at each serial position (**Fig.S4A**). We then asked 6 questions, as in (Konecky et al., 2017): can the first item be decoded in the first delay, in the second delay, or in the third delay; can the second item be decoded in the second or third delay; can the last item be decoded in the third delay (**Fig.S4B**)? The feature neurons strongly encoded the most-recently-presented item. Using the activity of conjunction units, however, nothing could be decoded above chance using a linear classifier.

#### Simulation 12: Previous trial repetition effect

This effect was examined by using all trials taken from simulation 1. We compared trials in which the probed-item’s features on the probe dimension and the recall dimension were the same or different to the probed item of the previous trial. Trials were grouped according to whether the previous trial’s probe color was the same as the current trial’s probe color, and also whether the probed-item’s orientation was the same or different to the previous trial’s probed-item orientation (**Fig.S4**). When the identical item was probed, recall was more accurate.

#### Simulation 13: Pattern similarity after intervening conjunctions

We ran 2000 trials of the 1-item memory condition, and examined delay-period activity in conjunction units. If neurons have classical receptive fields, then when the same stimulus is presented as on a previous trial, the activity pattern will be similar, whereas if the stimulus is different, the patterns will be dissimilar. Two major predictions of flexibly-conjunctive neurons is that the pattern similarity will decrease both with the number of intervening trials ( **Fig.5A**), and also when intervening stimuli form different conjunctions with the same features ( **Fig.5B-D**).

The similarity between representations on trial *n* and trials *n*-2, *n*-3 up to *n*-10 were examined, in the middle 100 timesteps of the delay period. For each inter-trial distance, trials were divided according to whether the same or different stimulus was shown on those two trials ( **Fig.5A**). We considered only 2 feature dimensions in this analysis, so that the each pair of trials had the same stimulus 25% of the time. For *n*-1 to *n*-4, it was possible to further subdivide trials according to the stimuli presented on intervening trials. A “violation” of the conjunction was defined as an intervening stimulus which is similar on one feature dimension but dissimilar on the other feature dimension, to trial *n*. No violation occurs if the intervening stimulus is the same as on trial *n*, or if it contains no features in common with trial *n*. Trials were split according to the number of these intervening violations.

#### Simulation 14: Implementing simple task rules

To implement execution of actions, we re-labelled the ‘orientation’ dimension as ‘action’, so that there were 4 possible motor actions, which could be coupled to the 4 possible colors ( **Fig.S6**). To provide the model with instructions for the task rules, each stimulus-response (S-R) mapping was activated, one at a time. For example, to provide the instruction “red means press button 1”, we activated the “red” colour unit and the “button 1” action unit simultaneously. Analogous to working memory encoding, a conjunction unit became associated with each pairing. In each block, we presented either 1, 2, 3 or 4 rules. After presenting the rules, 12 color cues randomly drawn from those rules were shown sequentially, analogous to memory probes. After each cue, activation of the motor units was measured. We tested the responses to one, two, three and four simultaneous S-R mappings.

The sequence of events was thus very similar to the working memory task, and reaction times and accuracy were calculated in the same way as before (**Fig.S6A**). Two differences in implementation were needed to permit appropriate action selection over many trials. First, we lengthened encoding (240 steps) and shortened the probe duration (80 steps). This increased the stability of the mappings. Second, rather than inhibiting the features during the inter-trial interval, no input was provided (**i** = **0** rather than **-1**), which allowed the units to maintain their ongoing activity. The attentional focus thus remained active throughout the experiment. Without these two changes, interference led to forgetting of the task rules over the first 5 to 10 trials.

Two important empirically observed effects in choice reaction time experiments are Hick’s law – the increase in RT with log(number of options) – and the repetition effect. To examine Hick’s law, we calculated the mean RT on correct trials, in blocks where there were 1 to 4 rules presented (**Fig.S6B**). To study the effect of stimulus-response repetition on consecutive trials, trials in the 2-rule blocks were categorized according to whether the same stimulus was present on the previous 0, 1 or 2 trials (**Fig.S6C**), and the mean RT was calculated separately for correct and incorrect response trials. Data for correct trials from (Schvaneveldt and Chase, 1969) were re-plotted next to simulations. Decoding from conjunctive units was performed as previously done for feature units, as a function of time. For each trial the classifier was trained on other trials in the same block (**Fig.S6D**). For comparison, accuracy when the classifier was trained on the previous block was also measured. During WM tasks no decoding was possible from conjunctive units. But in this simple stimulus-response task, decoding was possible across a block of trials which shared the same rule. The presentation of new task rules at the start of a block effectively overwrites the conjunctive neurons’ weights, precluding decoding.

#### Simulation 15: Removing the third (task-irrelevant) feature dimension

In previous simulations, each object consisted of three features, but one of the features is task-irrelevant. Simulation 3 was run with standard conditions and timings but on half of trials, the third dimension features were treated as absent (**i**=-1) for every object. We contrasted recall accuracy for conditions where objects consisted of all 3 features vs. only 2 features ( **Fig.S7**). The primacy and recency effects were observed to be smaller.

#### Simulation 16: Exploring the parameter space

We examined 9 free parameters in the model: the baseline activity, lateral inhibition × 2 (for conjunction and feature units), self-excitation × 2, reciprocal excitation × 2, and learning rates × 2 (**Fig.S8**). For each combination of parameters, 50 trials per condition were simulated for set-sizes 1 to 4, for all serial positions (500 trials per parameter set). The canonical model was perturbed along two parameters at a time. For each pair of parameters, a range of values above and below the canonical parameter values were tested, with 10 linearly-spaced levels of each parameter, to give a 10x10 grid. The parameter values used in Simulation 1 thus lie at the center of each grid, and the figure represents all possible cardinal planes through that point in a 9-dimensional hyperspace.

Each simulation resulted in a serial position curve like Fig.2D, from which we could quantify the set size effect (linear slope of accuracy as set size varies, collapsed across serial position), primacy effect (difference in accuracy between final and penultimate items in sequence, averaged across set sizes 2-4) and recency effect (difference in accuracy between first and second items in sequence, averaged across set sizes 2-4). The size of each effect was smoothed using a 3×3 boxcar, and is portrayed by pixel color in the 10x10 grids.

Minimal algorithm to reproduce main results

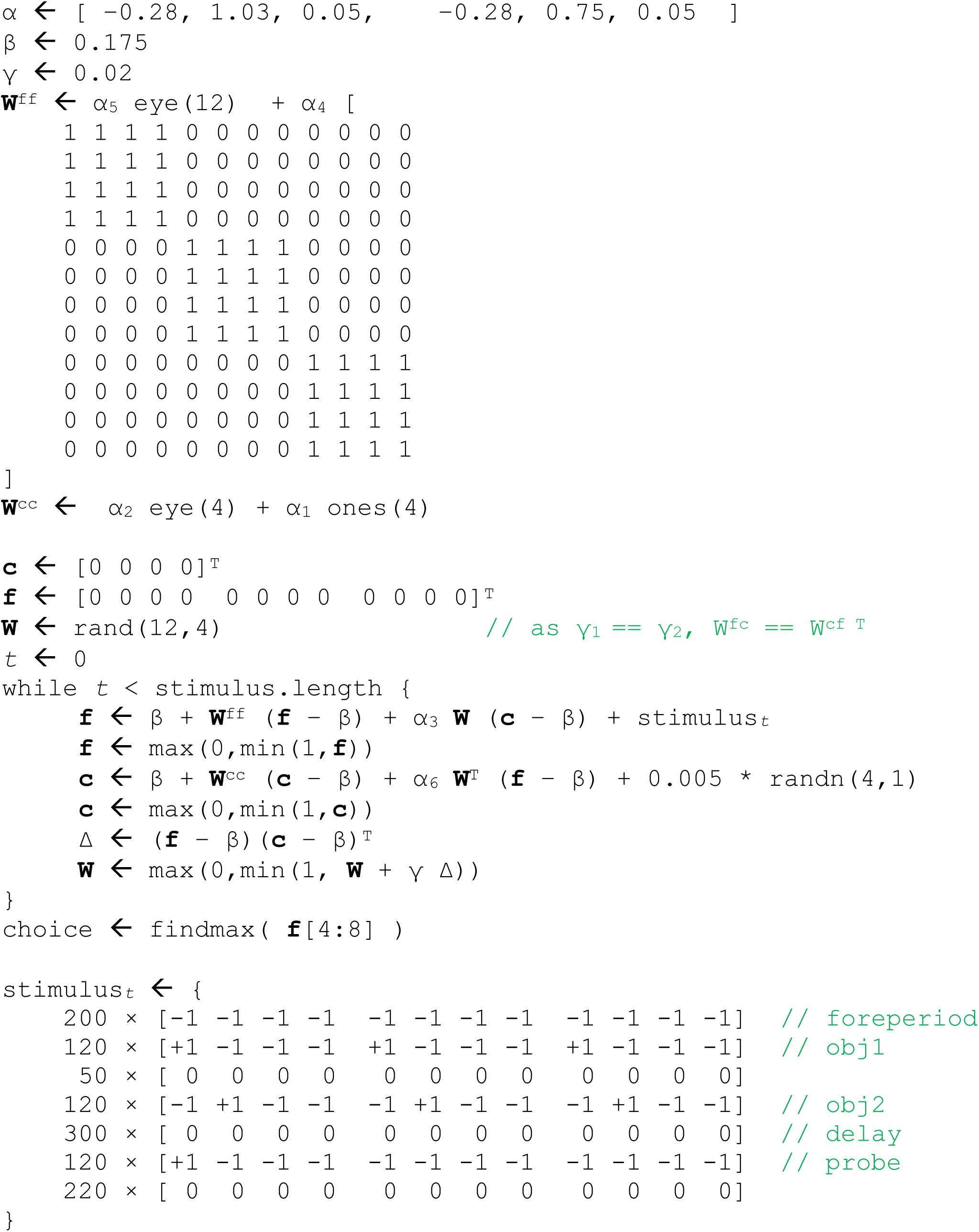

### Supplementary Figures

**Figure S1:**
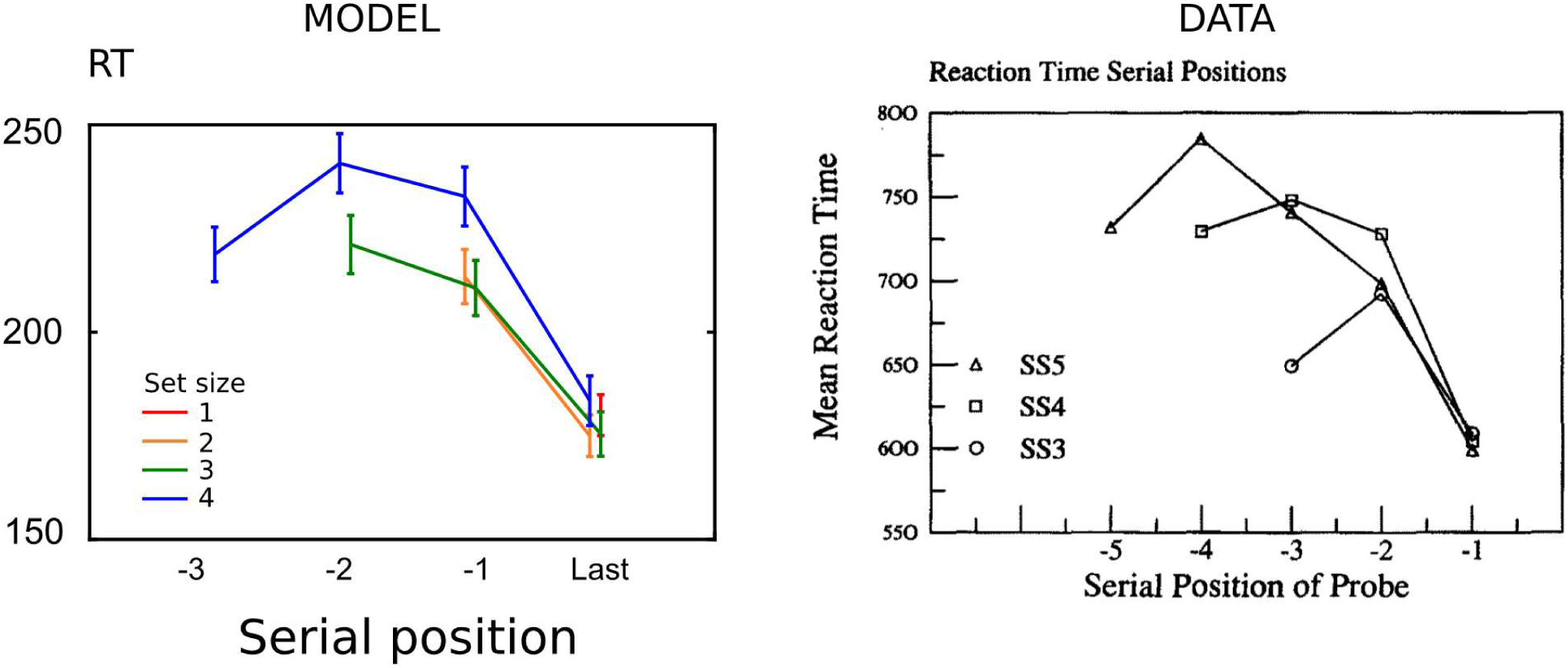
Reaction times (Simulation 7) The time taken to recall an item can be indexed by how fast the winning feature rises after probe presentation. An inverse relation with accuracy is observed, in keeping with behavior, with items already in the focus of attention showing the fastest responses. (**Simulation 8** below) A) Reaction times were extracted from trials of simulation 1, quantified as the time after the probe at which the winning feature reached 95% of its maximum value. The faster the attractor settled into a stable winning state, the shorter the reaction time. Each line represents a different set size, and the X-axis indicates serial position of the probed item within the memory set. Reaction times were approximately inversely related to accuracy on each condition. B) This is consistent with race-models of WM recall (Pearson et al., 2014), and aligns with behavioural data (McElree, 2006; McElree and Dosher, 1989)(Figure adapted from McElree & Dosher 1989).

**Figure S2:**
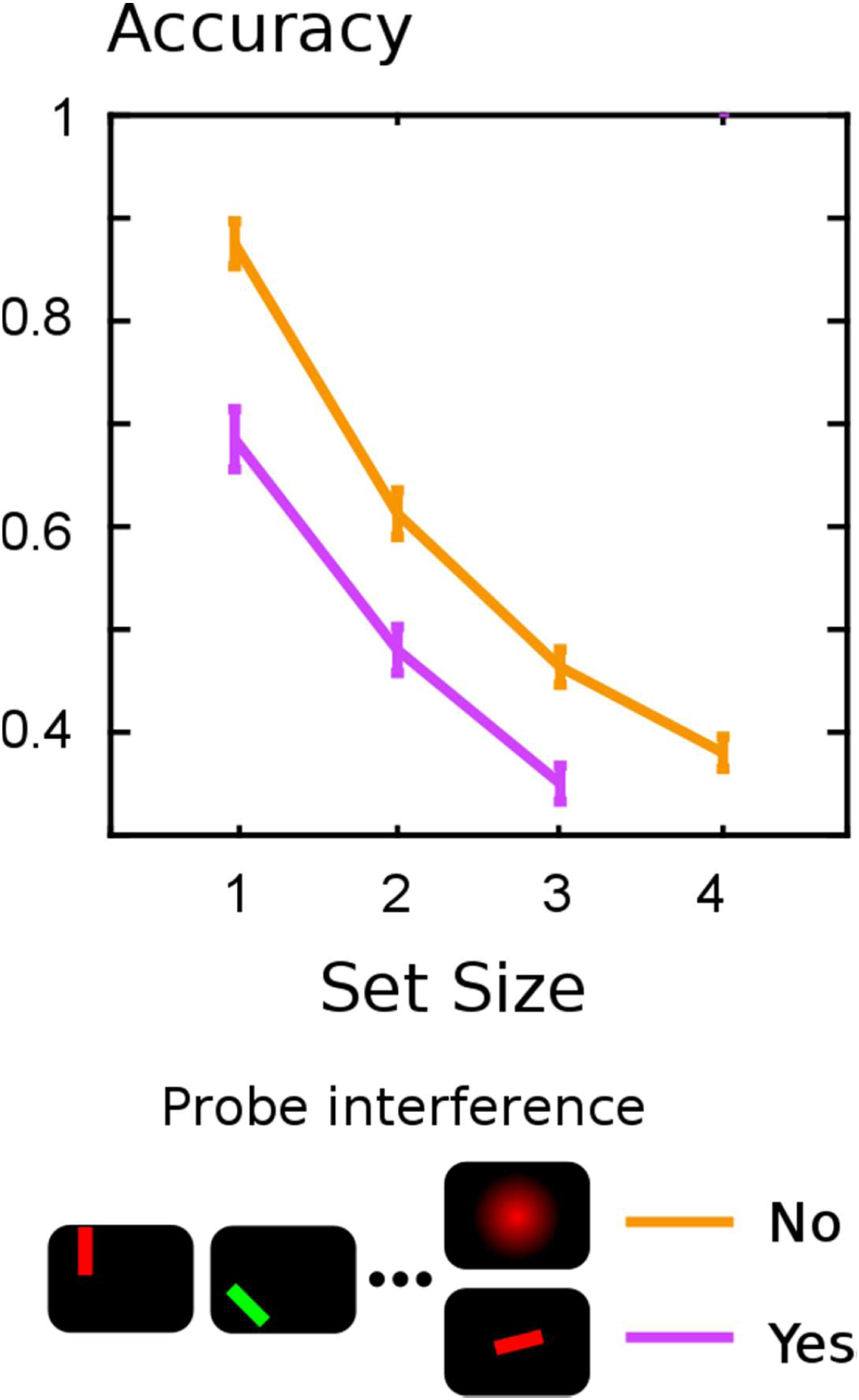
Novel prediction – Probe interference (Simulation 8) The model predicts that the probe can itself interfere with recall if it contains irrelevant features. The irrelevant features compete with the probe feature, and reduce the probability of correct refocusing of the item.

**Figure S3:**
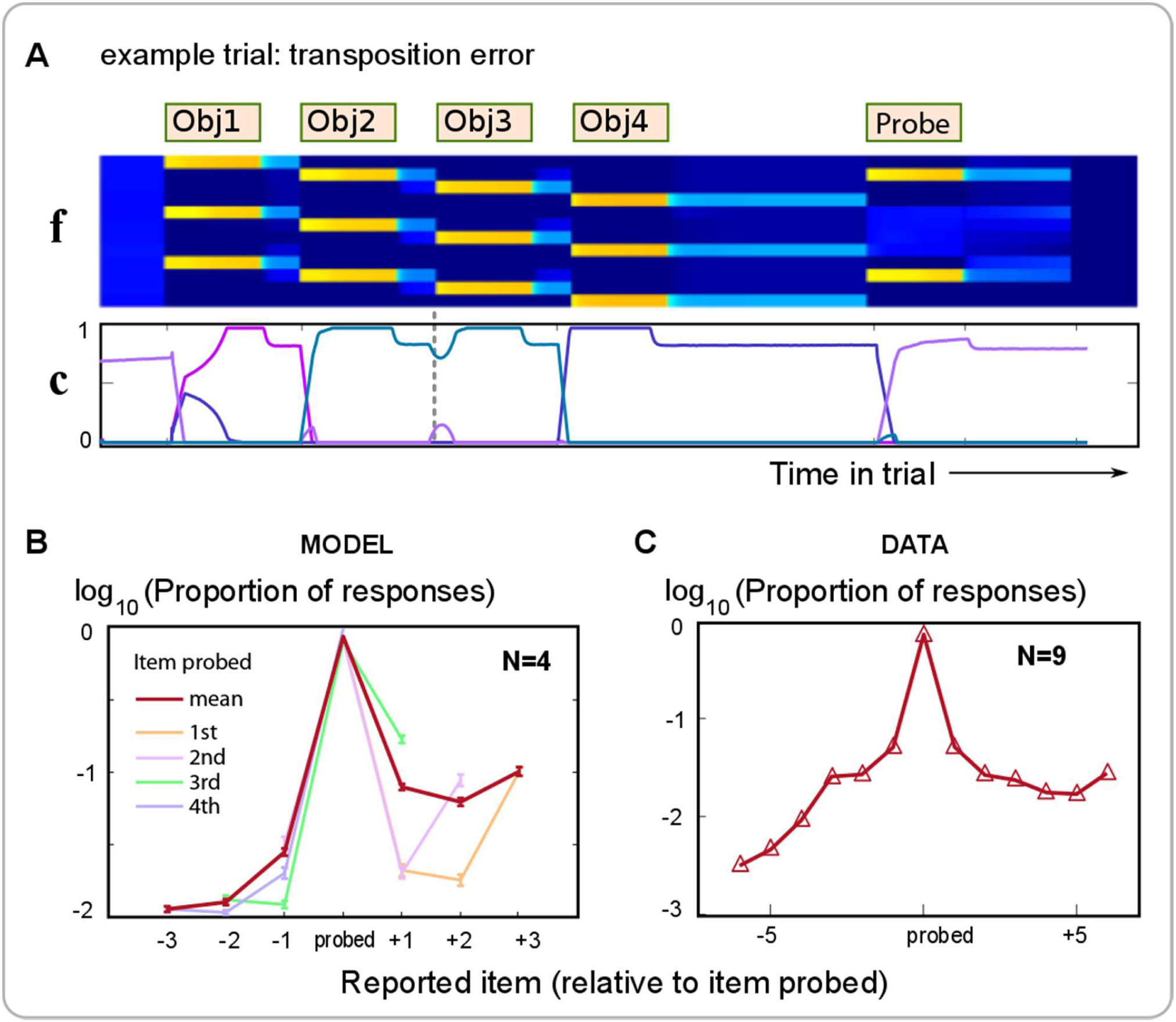
Errors are more likely to be sequentially neighbouring items (Simulation 5) **A** Transposition errors (sequence positional errors, or swap errors) arise when features of a temporally adjacent item are instead reported. In this example trial, the conjunction neuron activated by object 2 remained active for object 3 (vertical dashed line), so when item 2 or 3 is probed, the incorrect response feature is sometimes activated. **B** Items that were either just before (x<0), or just after (x>0), the probed item tended to be reported more often. **C** A qualitatively similar pattern of errors is observed in a verbal WM task in which a list of words had to be recalled in order (adapted from Expt.3 of Farrell and Lewandowsky, 2004). A very similar falloff with inter-item distance is also seen in visuospatial memory using sequence change-detection (Smyth and Scholey, 1996), but their data did not permit distinguishing forward and backward transpositions.

**Figure S4:**
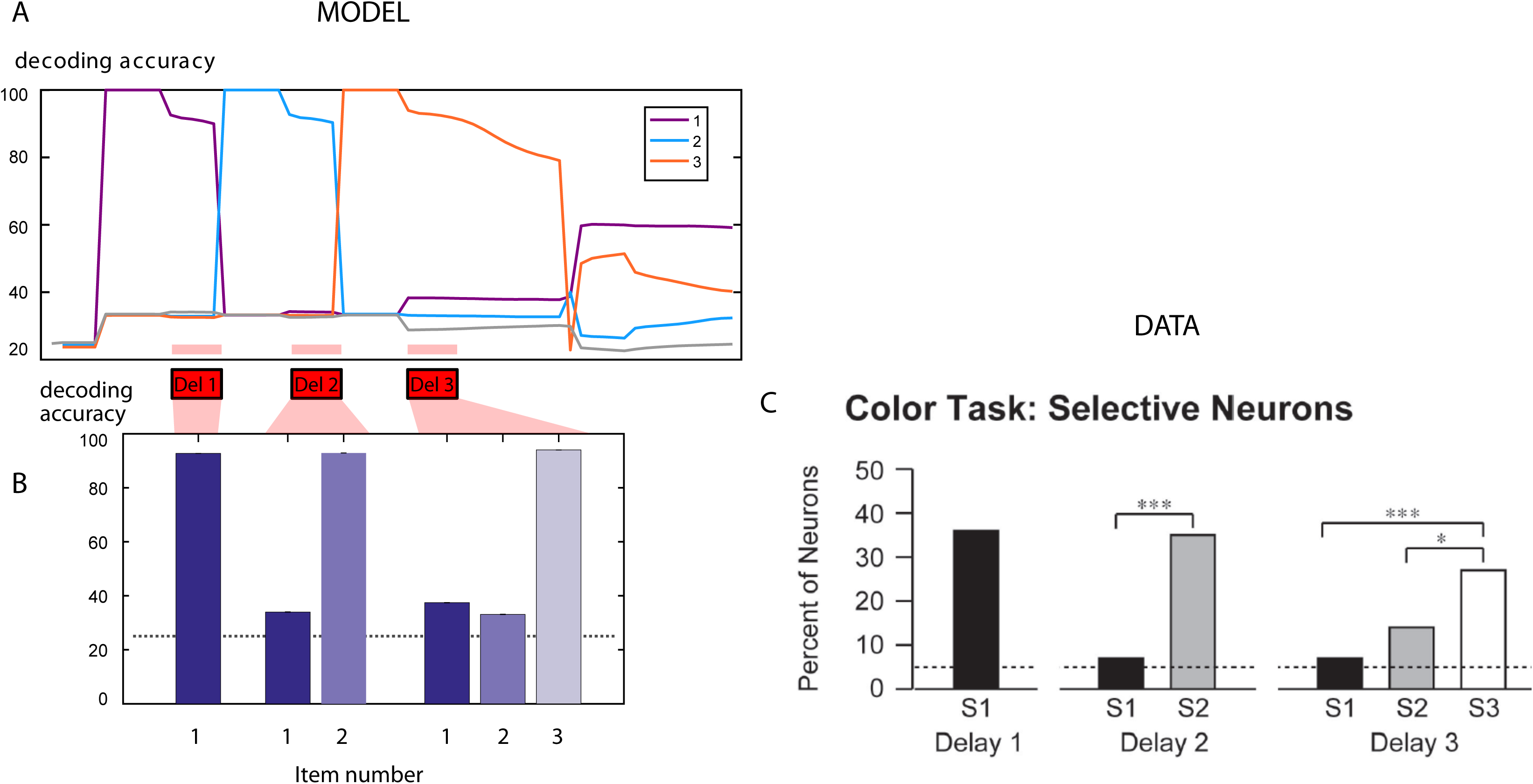
Decoding from feature units during the delay period (Simulation 11) Feature-neuron activity during the delay period of the 3-item condition was examined. For each time point, the activity of the four color units was used to decode (across trials) the identity of each presented color. A linear classifier was trained on the activity in the feature units **f** on 50% of trials, and tested on the remaining trials. A) The decoder accuracy for each of the three items is shown, as a function of time. The three delay periods of interest, following the presentation of each item, are shown as pink bars below. B) Average decoder accuracy during each of the delays. During the first delay we could decode the identity of the first item’s colour. During the second delay, we could decoded the second items’ color but not the first item’s color. In the third delay we could decode the third item’s color but not the other two colors. Dotted line is chance. C) Data from (Konecky et al., 2017), showing decodability of only the most recent item presented during a monkey WM task. The neurons were in fact from the principal sulcus, and 35% of neurons recorded here were feature-selective, though the remaining neurons were not.

**Figure S5:**
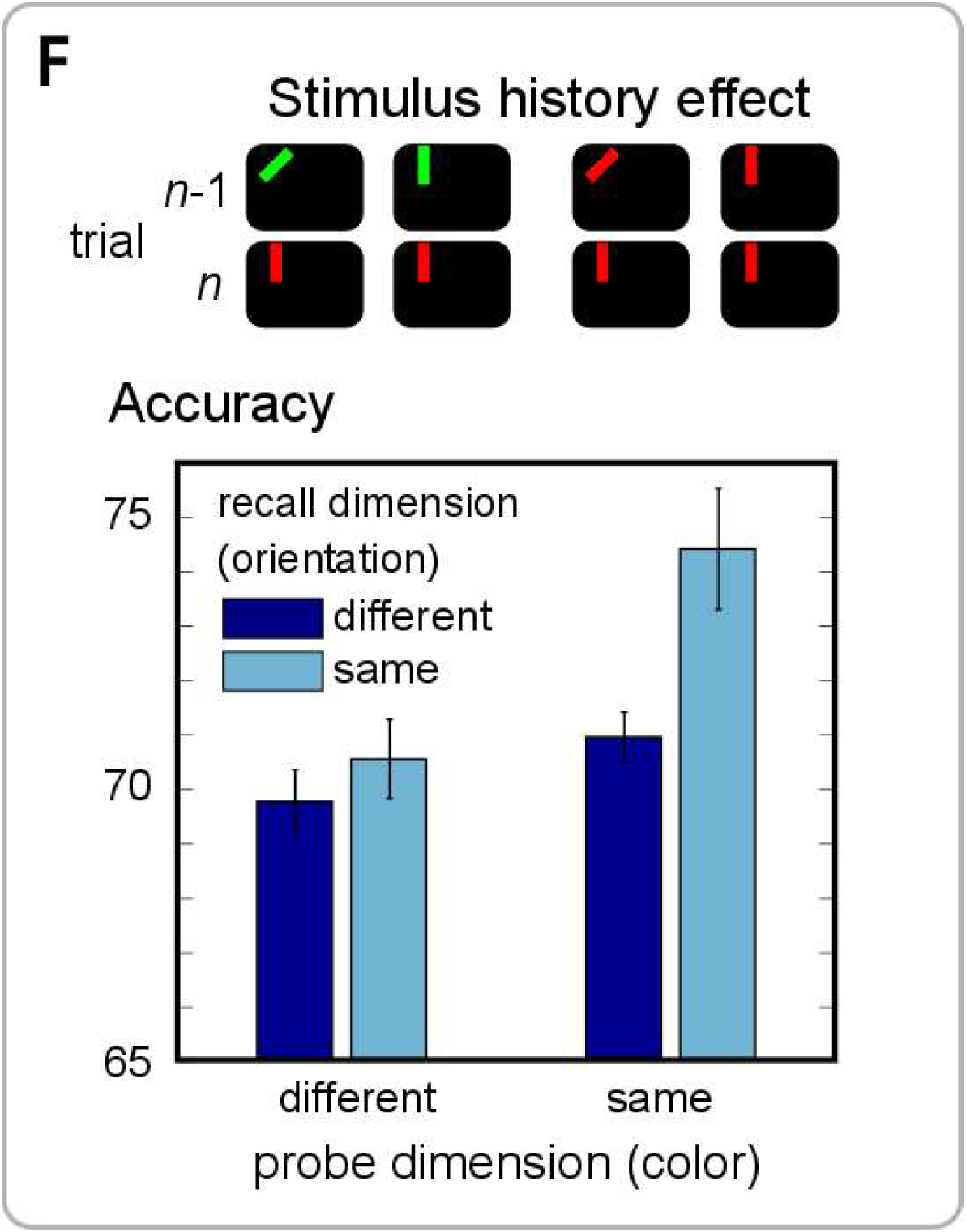
Novel prediction: Trial-to-trial conjunctions effect (Simulation 12) Since conjunctive neuron selectivities rely upon synaptic traces from the trial history, we predict that the stimuli presented on the previous trial generate interference effects with the current trial. The model predicted that when the probed item’s features were identical to those of the item probed on the previous trial, responses were more accurate.

**Figure S6:**
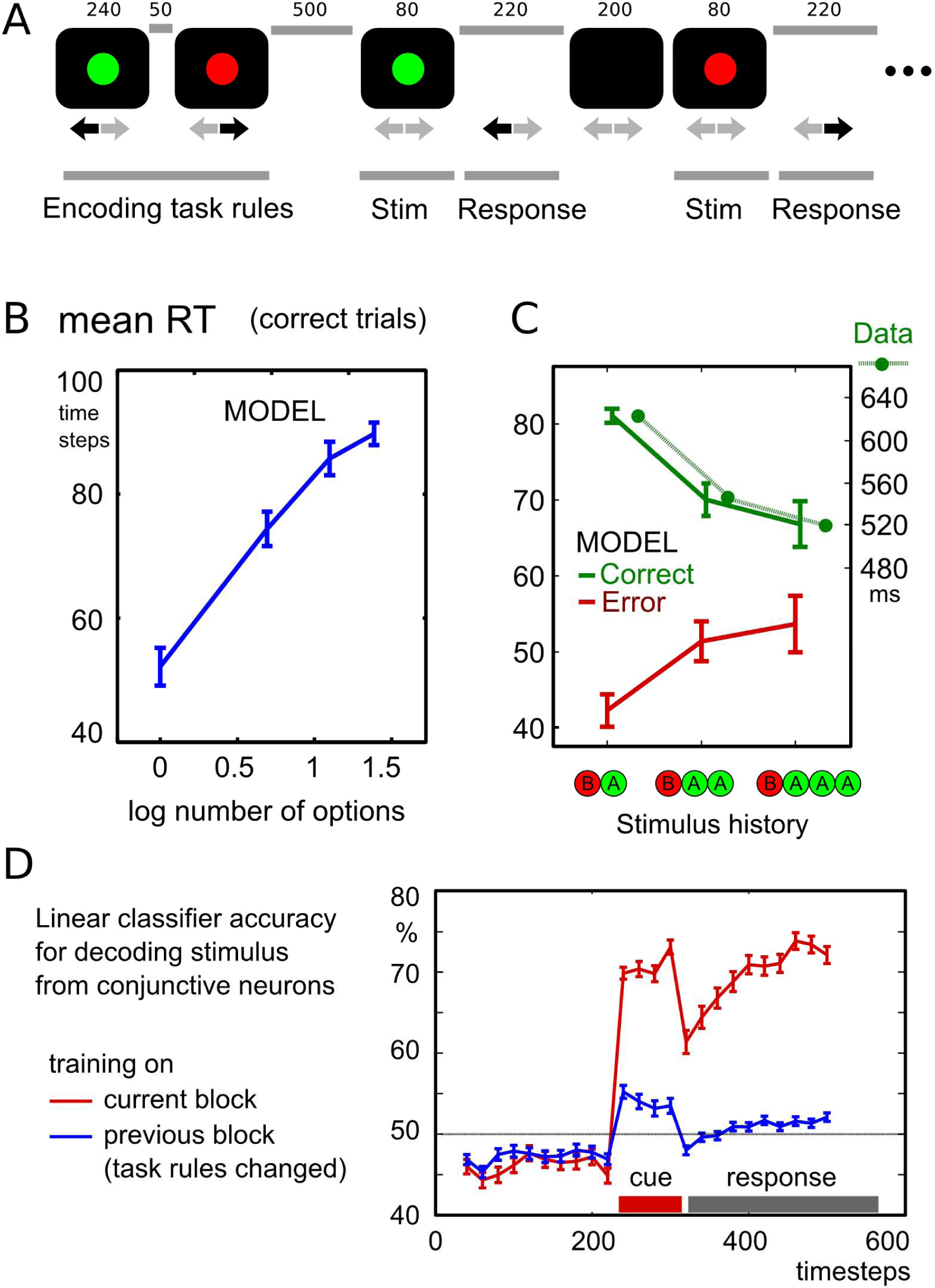
Acting on multiple task rules (Simulation 14) Here we simulated a simple N-alternative choice task, where each colour indicates that a particular button should be pressed. A) At the start of each block, a set of task rules was encoded. A single color and motor plan feature were activated simultaneously, indicating the rule for example “if red, press left”. After 1 to 4 rules were presented, a series of test trials followed. One of the colour features that was presented previously was activated, corresponding to presenting a cue. The subsequent response unit activation was measured, to indicate the model’s response. Reaction times were calculated as previously from the time of stimulus onset. B) The RT increased with the logarithm of the number of rules encoded, according to Hick’s law. C) RTs were split according to stimulus repetition history. If the same stimulus was tested on the previous trial (‘BAA’), or on the previous two trials (‘BAAA’), then the RT on correct trials was faster than if the stimulus was different (‘BA’), in keeping with data. Dotted line: RT on correct trials replotted from Expt 4 of (Schvaneveldt and Chase, 1969), in which one of 4 responses was selected after seeing one of 4 stimuli, according to an arbitrary stimulus-response mapping. The model also predicts errors will be faster than correct responses, with an inverted stimulus-repetition effect. D) Unlike in the WM task, decoding is possible from conjunctive units, since the task set is maintained rather than overwritten on each trial. For each test trial, the stimulus identity was decoded using the other trials in the same block for training (red), or the trials in the previous block (blue). Decoding was possible within the block, because a consistent conjunction unit pattern – representing the task rule corresponding to the current stimulus – was activated.

**Fig. S7:**
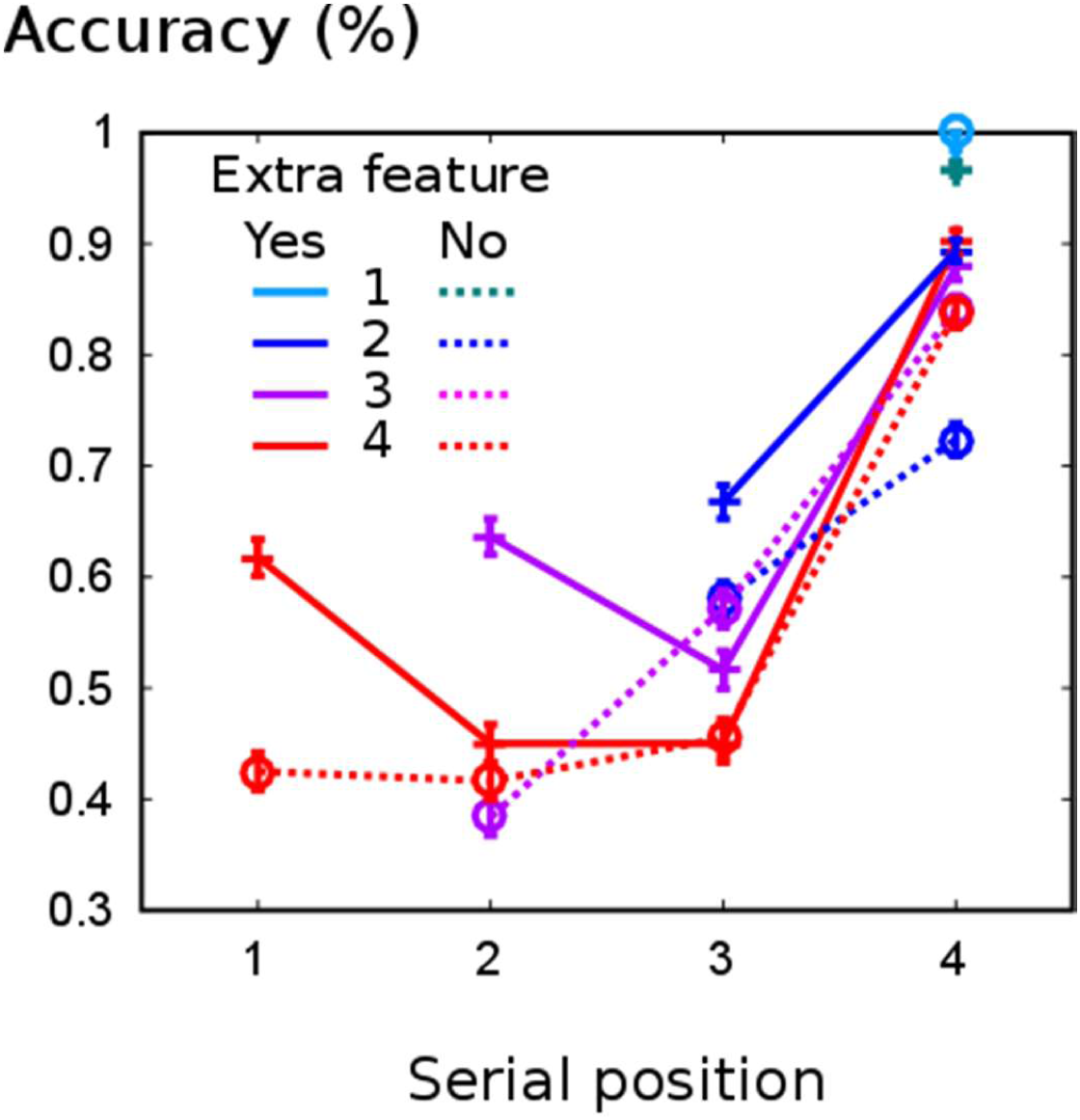
Novel prediction: Benefit with an extra feature dimension (Simulation 15) Previous simulations had three features per object (colour, orientation and location), of which only two were task-relevant. The third, irrelevant, feature was distinct for each object. Removing the task-irrelevant feature (e.g., presenting sequential items all at one location, rather than at different locations) worsens model performance, in particular by reducing primacy and recency benefits.

**Fig.S8:**
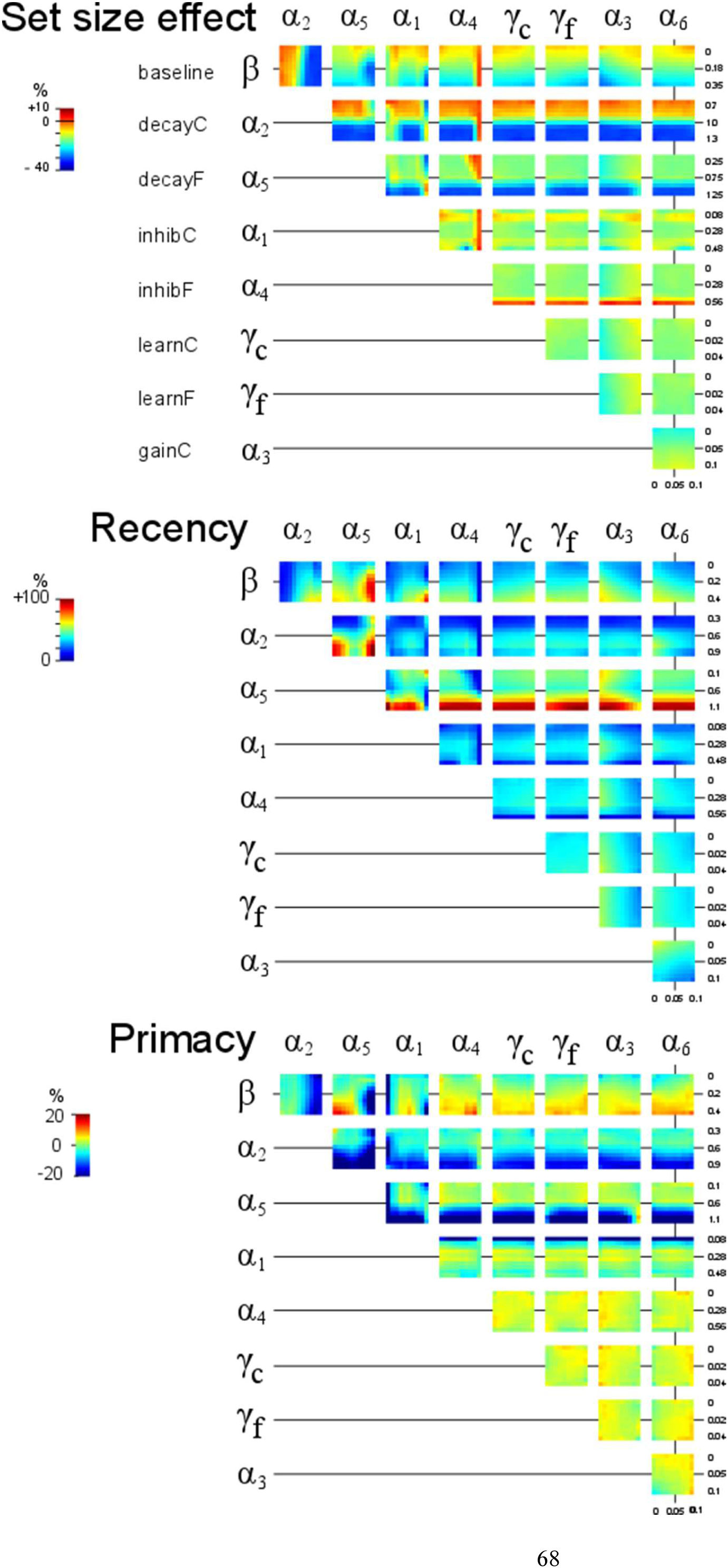
Influence of model parameters upon central behavioral effects (Simulation 16) We examined the effect of varying 9 free parameters in the model, varying two of them at a time. The parameter values used in the main paper lie at the centre of each 10 × 10 grid, and the figure represents all possible pairs of free parameters. For each parameter combination, we quantified the set size effect (reduction in accuracy as set size increases), primacy effect (difference in accuracy between final and penultimate items in sequence) and recency effect (difference in accuracy between first and second items in sequence. Warm pixels indicate larger effects.

**Fig.S9:**
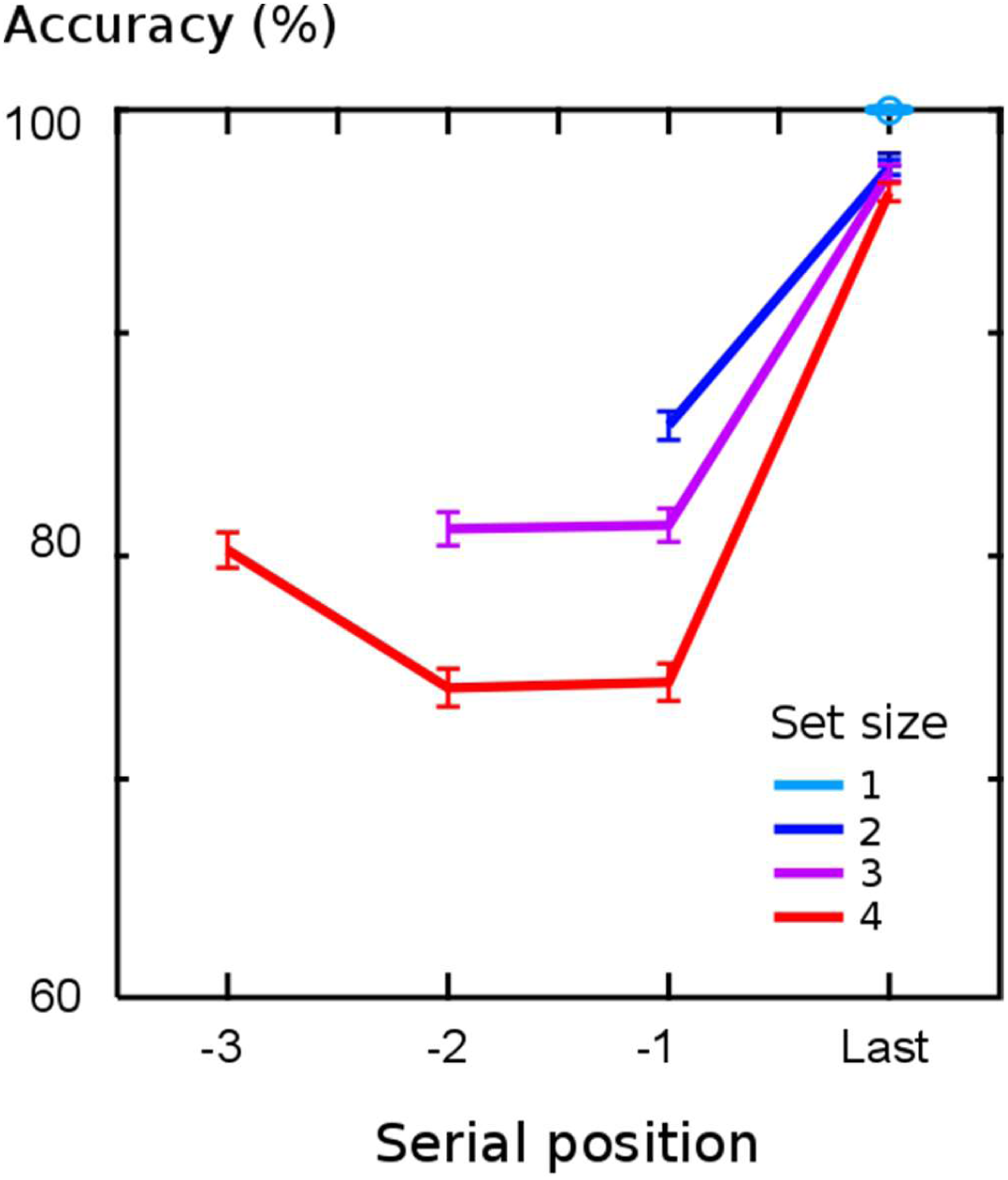
Basic results for ‘high-accuracy’ parameter regime (used for simulations 3 to 5) To prevent floor effects when simulating encoding and delay effects, we required a higher initial performance level, and adjusted the parameters accordingly. This figure demonstrates the equivalent of Figure 2C, using this new set of parameters. Performance is overall higher but demonstrates qualitatively similar effects.

**Table S1:** Empirical findings explained by the model

| **Phenomenon explained** | **Mechanism in model** | **Refs** |
| --- | --- | --- |
| **Set size** reduces recall accuracy | Competition between conjunction units → interference | (Zhang and Luck, 2008) |
| **Primacy** – first item benefit | No competition with previous items at encoding | (Baddeley, 1996) |
| **Recency** – last item benefit | Features retained in active state; no retrograde interference | (Gorgoraptis et al., 2011) |
| Only last item **encoded actively** in firing rates | Subsequent items capture focus of attention | (Konecky et al., 2017) |
| **Shifting attention** to item improves recall | Re-activation by pattern completion focuses attention on item by sustained activity. Subsequently probing that item is faster and more accurate, as it is already active. | (Myers et al., 2017; Souza et al., 2016; Zokaei et al., 2014b) |
| **Additional features** in an object remembered without extra cost | During encoding, synaptic weights increase in parallel to all concurrently active features. | (Allen et al., 2006; Luck and Vogel, 1997; Sala and Courtney, 2007) |
| **Frontoparietal activation** during WM maintenance | Conjunctive and feature neuron firing required for focus of attention and thus for encoding | (Rowe et al., 2000; Sakai et al., 2002) |
| **Frontoparietal connectivity** is signature of shifting attention | Synapses between conjunctive and feature units are bidirectionally activated to drive persistent activity | (Heinen et al., 2017; Scolari et al., 2015; Szczepanski et al., 2013) |
| **Synchrony** between posterior and frontal regions during WM | Reciprocal excitation between conjunctive and feature neurons crucial for stable attractor of focus of attention. | (Fries et al., 2001; Gregoriou et al., 2009) |
| Fine-grained decoding from PFC during WM is elusive | Conjunctive neurons encode information flexibly across trials, with selectivity depending on recent history | (Cogan et al., 2017; Harrison and Tong, 2009; Lara and Wallis, 2014; Sprague et al., 2016) |
| Working memory operates as an **attentional template** | Unattended items in WM maintained synaptically so stable attractor is reactivated by partial information | (Lavie and Fockert, 2005; Woodman et al., 2007) |
| Memory contents lead to obligatory **capture of attention** | Partial information pertaining to items in silent WM is amplified, leading to persistent activity / focus of attention | (Olivers et al., 2006; Soto et al., 2008) |
| WM capacity correlates with **complexity of task set** | One WM object corresponds to one stimulus-response pairing | (Conway et al., 2003; Duncan, 2010) |
| PFC subserves both WM and task set maintenance | Conjunctive neurons can flexibly bind actions to stimulus features, or groups of features, together | (Barch et al., 1997; Duncan et al., 2000) |
| Neural decoding of **unattended items** in WM is weak | Unattended items encoded in synaptic traces | (Lewis-Peacock et al., 2012; Sprague et al., 2016) |
| TMS to feature areas, or indiscriminate sensory stimuli, can **re-activate** representations | Neurons reactivated by TMS pulse that are connected synaptically to conjunctive neurons (i.e. unattended WM items) are selectively amplified by reciprocal synapses | (Rose et al., 2016; Wolff et al., 2017) |
| **TMS** to posterior cortex disrupts benefit conferred by focus of attention | Electrical activity i.e. persistent activation is disrupted by electrical stimulus, but synapses are not | (Zokaei et al., 2014a) |
| **rTMS to PFC** disrupts WM by altering posterior activity | Reducing conjunctive unit excitability reduces stability of attentional attractors | (Zanto et al., 2011) |
| Attention operates by frontal **amplification** of posterior feature-selective neurons | The attractor basin formed by mutual conjunctive and feature synapses allows conjunctive units to control gain in feature-selective neurons. | (Desimone and Duncan, 1995; Merrikhi et al., 2017; Moore and Armstrong, 2003) |
| Primacy effect not robust | Depends on ITI and events before trial | (BADDELEY, 2000) |
| **Transposition** errors | New conjunction unit not always activated for new items | (Farrell and Lewandowsky, 2004) |
| Binding must be **serial** | Conjunctive neurons encode all simultaneously active features. | (McLean et al., 1983; Treisman and Gelade, 1980) |
| More items mean faster decay | Focus of attention is robust to decay | (Pertsov et al., 2016) |
| Errors report succeeding items more than prior items | More recently encoded items have stronger synaptic weights, and more likely to intrude. | (Farrell and Lewandowsky, 2004) |
| **Probe can interfere** with recall | Irrelevant features in probe suppress reactivation of items | (Souza et al., 2016) |
| **Reaction times** inversely related to accuracy | Time taken for conjunctive unit to activate features depends on synaptic strength and competition from other units | (McElree and Doshier, 1989; Pearson et al., 2014) |
| **Encoding rate** slower when more items stored | Items encoded briefly are more susceptible to interference from other items in memory | (Bays et al., 2011) |

**Table S2:** Psychological concepts corresponding to the model

|  |  |
| --- | --- |
| **Binding** | Simultaneous activation of two feature neurons causes a single conjunction neuron to become active and form bidirectional connections to those feature neurons. |
| **Focus of attention** | Active representation in feature-selective neurons, coupled with a conjunctive unit that is simultaneously active. The two types of neuron are mutually excitatory and generate persistent activity. |
| **Unfocused item in WM** | Bidirectional increases in synaptic weights between a combination of place-coded feature neurons, and one conjunction neuron |
| **Recall** | Associative re-activation of a pattern of activity in feature neurons. |
| **Task set** | One of the feature dimensions represents motor-plan neurons. During encoding, the simultaneous activation of a motor plan and a perceptual feature causes a conjunctive unit to associate the two. Reactivating the perceptual feature triggers the motor program. |
| **Top-down control** | Conjunction neurons effectively amplify feature neurons that were previously encoded as WM items. |
| **Forgetting from WM** | Activation of feature neurons by an external stimulus provides new input. This input competes to be represented by conjunction neurons. The conjunction neuron with the most-similar connections will be activated and re-wire to encode the stimulus. This interfering stimulus thus overwrites and displaces an item previously in WM. |

**Movie S1**: **Timecourse of activity during working memory encoding and recall**. (A) top left panel shows the instantaneous rate of change in weights Δ, (B) below is shown the current synaptic weights. (C) top right illustrates the object currently encoded by the feature neurons. The drawn intensity of each possible stimulus is the product of the activity of the corresponding feature neurons. During encoding this corresponds to the objects presented to the model. (D) lower panels show the feature neuron activity as a heatmap, as a function of time, and the activities of the four conjunctive neurons as traces. Delay period activity generally corresponds to the final item presented. Errors occur when one conjunctive neuron is active for two objects, or when one conjunctive unit fails to win the competition.

